# Tissue-specific chromatin accessibility and co-factor availability together define ASCL1-dependent neural reprogrammability across germ layers during embryogenesis

**DOI:** 10.64898/2026.07.13.738233

**Authors:** David Lando, Samina Kausar, Toshiaki Shigeoka, Frances Connor, Jerome Jullien, Anna Philpott

## Abstract

The proneural transcription factor ASCL1 is well established as a pioneering neuronal reprogramming factor in several contexts. However, the extent to which distinct tissue types adopt a similar or divergent response to the same ASCL1 fate challenge is unclear. By expressing ASCL1 across germ layers and tissues of the developing *Xenopus* neurula embryo, we reveal widespread but largely distinct transcriptomic and chromatin responses to ASCL1. Ectopic ASCL1 can access partially open chromatin sites across all tissues, but it is able to further open those sites specifically in neuroectoderm. Using motif enrichment and accessibility analyses we identify Sox transcription factor motifs as enriched and preferentially accessible at ASCL1-bound regions in neuroectoderm. We show that co-expression of Sox3 and ASCL1 can enable the activation of otherwise refractory ASCL1 target genes in mesoderm and epidermal skin. Our findings reveal how the chromatin landscape and co-factor availability work together to modulate the response to transcription factor-driven fate challenge during development.

**Summary statement:** Single-cell multi-omics reveals how chromatin accessibility and cofactor availability constrains ASCL1-driven neuronal reprogramming across tissues in the developing *Xenopus* embryo.

## Introduction

Development of an organism depends on the precise regulation of cell fate diversification, yet the mechanisms that control cellular competence for correct lineage specification at the tissue level are not fully resolved. Compared to early development in the mammalian embryo, *Xenopus* frog embryos provide a more tractable and accessible system for understanding the gene regulatory mechanisms that control cellular competence; moreover, these mechanisms are usually found to be conserved across species (Sive, 2023).

Developmental programmes in *Xenopus* are initiated by maternally deposited transcription factors which activate zygotic gene expression. This drives the differentiation of pluripotent cells into the three primary germ layers, ectoderm, mesoderm, and endoderm, followed by the emergence of organ progenitors (Paraiso et al., 2020, Zhou and Cho, 2022). The onset of germ layer specification coincides with extensive reorganisation of the embryonic chromatin landscape, including dynamic changes in chromatin accessibility and histone modifications that contribute to cell fate acquisition. Chromatin accessibility is characterized by a dynamic transition from a mostly closed state in early embryonic cleavage stages to a highly accessible state at the blastula stage when transcriptional activation of the zygotic genome occurs (Esmaeili et al., 2020, Cui et al., 2025). This opening is mediated in part by pioneer transcription factors from the maternal oocyte that bind to closed chromatin and remodel it into a more accessible state (Cui et al., 2025). Once opened gene promoters become marked by histone H3K4me3, while cis-regulatory active enhancers are marked by H3K27ac (Gupta et al., 2014, Hontelez et al., 2015, Akkers et al., 2009). The gradual loss of cell competence to respond to early developmental signals has been linked to the closing of chromatin at specific promoters. For example, in Wnt signalling, Wnt target genes on the dorsal side of the embryo, such as *nodal3.1,* become inaccessible as embryos begin gastrulation (Esmaeili et al., 2020). Hence, the status of chromatin accessibility at gene promoters and likely also at cis-regulatory enhancers helps to shape transcriptional competence, thereby constraining lineage choice during early *Xenopus* embryogenesis.

Lineage choice is typically directed by key transcription factor “master regulators”. One examples of these is ASCL1, a pro-neural basic helix–loop–helix (bHLH) transcription factor that functions as a master regulator of neural differentiation in Drosophila, (Lo et al., 1991, mammals (Guillemot et al., 1993) and Xenopus (Ferreiro et al., 1993). During *Xenopus* development, a role for maternally expressed ASCL1 has been implicated in defining the germ layers (Gao et al., 2016), Zygotically expressed ASCL1 is found in the anterior regions of the central nervous system (CNS) (Ferreiro et al., 1993) where it has been shown to drive neurogenesis (Ferreiro et al., 1993, Zimmerman et al., 1993, Ferreiro et al., 1994). Given its conserved role as a master regulator, ASCL1 has been used as a key component in a variety of methods to reprogram mammalian cells into neurons (Vierbuchen et al., 2010, Caiazzo et al., 2011, Son et al., 2011, Pfisterer et al., 2011).

Ectopic over-expression of ASCL1 in the developing *Xenopus* embryo leads to neuronal differentiation, as defined by the expression of neural markers such as neural beta-tubulin, primarily in dorsal ectoderm tissue; by contrast, ventral mesoderm and endoderm tissue remain resistant to reprogramming (Talikka et al., 2002, Ali et al., 2011, Hardwick and Philpott, 2018). This suggests that ASCL1-mediated reprogramming is highly influenced by cellular context, but what determines the competence of a cell to respond to ASCL1 in the *Xenopus* embryo is currently not clear. Some mammalian neuronal reprogramming approaches using ASCL1 have shown that it can act as a “pioneer factor”, binding and directly opening otherwise closed nucleosomal DNA in some non-neuronal cell types, such as fibroblasts (Wapinski et al., 2013, Wapinski et al., 2017, Zhou et al., 2026). However, other cell types, such as keratinocytes remain refractory to ASCL1-mediated reprogramming (Wapinski et al., 2013). The ability of ASCL1 to access and open chromatin only in specific cell types at specific stages of differentiation could explain the differing competences of tissues to respond to ASCL1-mediated reprogramming.

In this study, we investigate possible mechanisms that could determine the ability of different tissues in the developing Xenopus embryo to respond to the same ASCL1-mediated fate challenge. Specifically, we ask if the response of different tissues to ASCL1 is primarily controlled by chromatin accessibility and/or by the availability of cofactors that ASCL1 requires to direct a neuronal programme. We find evidence of both control mechanisms: where cell-type-specific responses to ASCL1-induced neuronal gene activation across different tissues in neurula embryos are driven by both the efficiency with which ASCL1 can further open chromatin at regulatory elements, as well as the tissue-specific availability of Sox co-factors that are needed for ASCL1 to access a neurogenic programme. Notably, embryonic tissues differ significantly in cofactor expression and motif accessibility near ASCL1 binding sites, which directly dictates their response to ASCL1.

## Results

### Ectopic ASCL1 activation in *Xenopus* neurula embryo results in tissue-specific transcriptomic and chromatin accessibility changes

Exploiting the advantages of the Xenopus system, we developed a method to explore the potential cell fate choices made by different embryonic tissues in response to similar ASCL1 activation in vivo at neurula stage, when germ layers are well established and tissue specification is already well underway (Fig. 1A). We injected fertilised Xenopus eggs with mRNA encoding full length mouse ASCL1 fused to the ligand binding domain of the glucocorticoid receptor (ASCL1-GR) (Talikka et al., 2002, Hollenberg et al., 1993) or GFP (WT). Embryos where then cultured to late neurula stage 18 when dexamethasone (Dex), a GR ligand, was added to activate ASCL1-GR that wass now found associate with chromatin (Fig. S1A). We then collected ASCL1-GR expressing embryos at three developmental timepoints, T0, T1.25 and T2.5 hours (hr) after the addition of dexamethasome, embryonic stages 18, 19/20 and 21 respectively, alongside control wild-type embryos injected with GFP taken at the same time-points. This sampling window captures the developing embryo as it undergoes rapid neural tube closure and neural regionalization (Tarin, 1971) and is designed to measure the response of different cell types at the same developmental stage to a similar ASCL1 reprogramming challenge.

**Figure 1.**
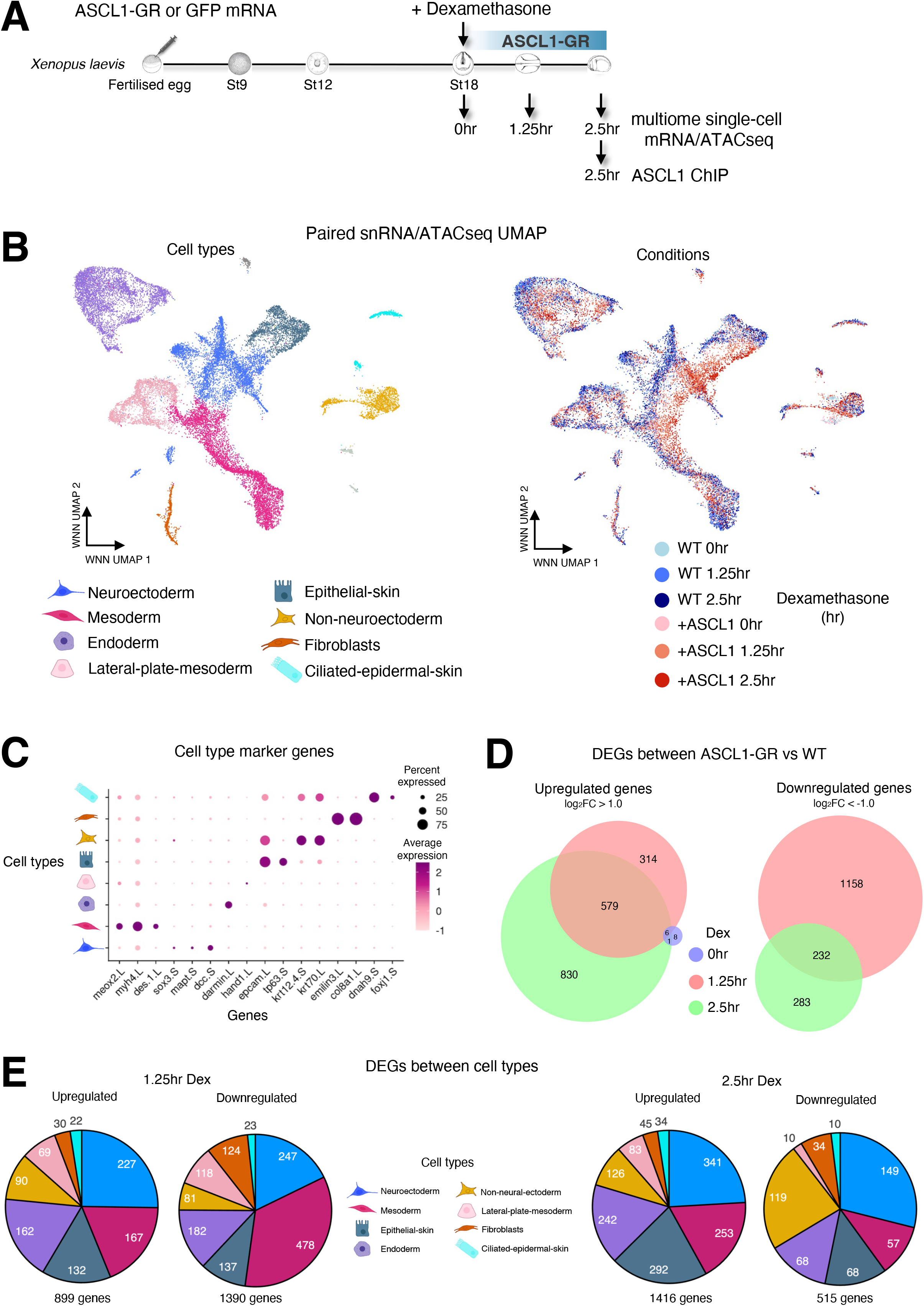
Study of ASCL1 directed reprograming in Xenopus development. A. Schematic of the approach and samples collected to investigate ASCL1 directed reprogramming in *Xenopus*. See main text for description. B. Weighted nearest neighbour (WNN) UMAP of 24,282 integrated single-cell multiome paired RNA and ATAC-seq (snmRNA/ATACseq) profiles for WT (GFP mRNA) and +ASCL1 (ASCL1-GR mRNA) samples from three dexamethasone treated time points, coloured by cell type annotation (left) or time point (right). C. Dotplot of the average expression of markers genes for each cell type annotation. D. Venn diagram plot of the total number of ASCL1-GR vs WT differentially expressed genes (DEGs) up regulated (left) and downregulated (right) between each cell type for the three timepoints after dexamethasone (Dex) treatment. E. Plot of the number of differentially expressed genes (DEGs) up and downregulated for each cell type from (D) after 1.25hr (left) and 2.5hr (right) dexamethasone (Dex) treatment.

**Figure S1.**
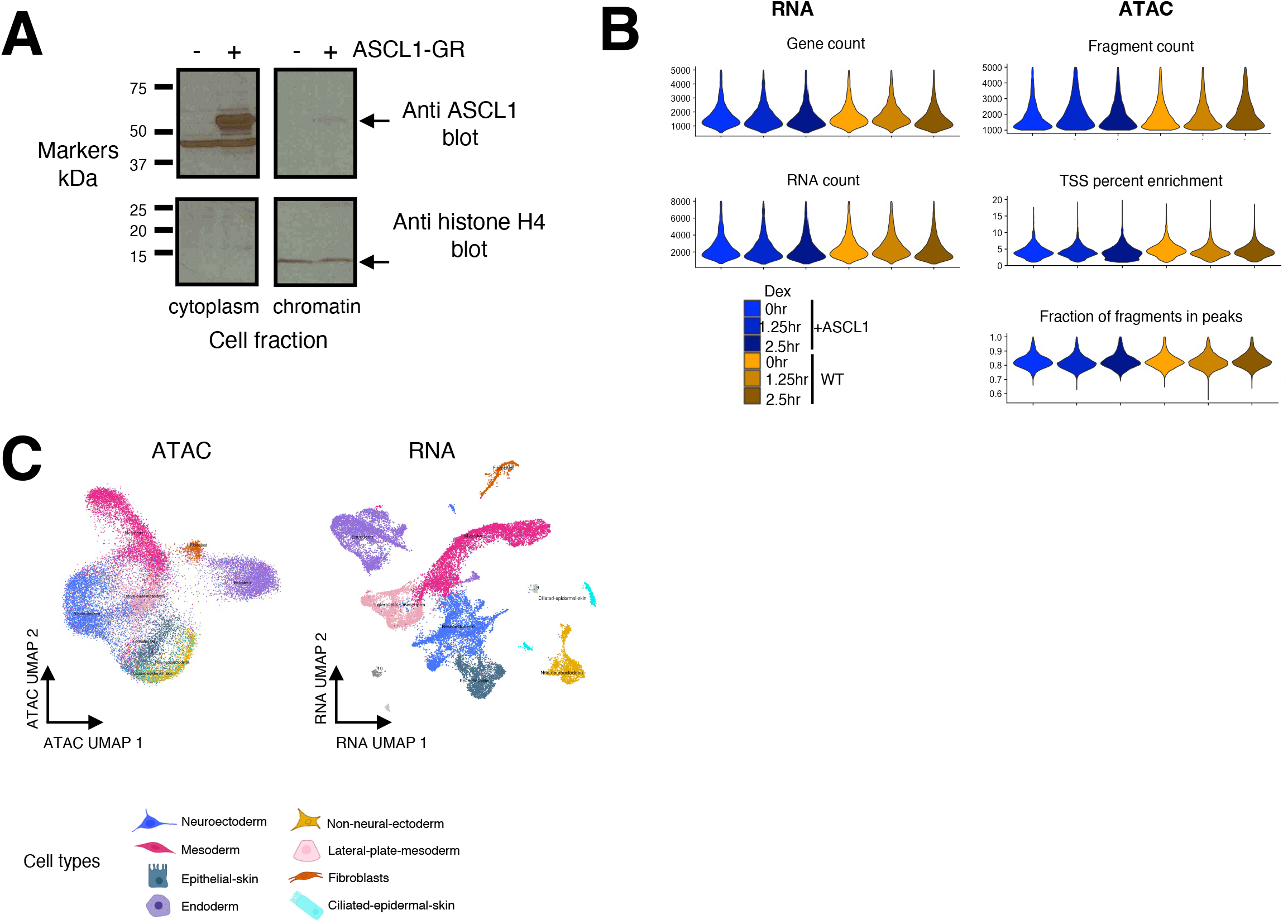
Quality control analysis of snRNA/ATAC-seq. A. Western blot analysis of cytoplasmic and chromatin fraction of GFP RNA (-) and ASCL1GR RNA (+) injected embryos at stage18. Top and bottom panels were blotted using anti rabbit polyclonal antibody to ASCL1 and histone H4 respectively. Arrow indicates the position of ASCL1-GR (top panel) and histone H4 (bottom panel). B. Violin plots of the gene count and RNA count for WT and +ASCL1 treated dexamethasone (Dex) samples for the single-cell RNA data (left panels) and single-cell ATAC data of the fragment count, transcriptional start site (TSS) enrichment and fraction of fragments in ATAC called peaks (right panels). C. UMAP profile of all single-cell multiome samples from Figure 1B split by ATAC (left panel) and RNA (right panel) and coloured by cell type annotation.

*Xenopus* embryos contain large numbers of yolk platelets that store vital nutrients for the developing embryo (Jorgensen et al., 2009), which can make isolation of cells and nuclei for single cell analysis challenging. To optimise single cell/nuclei analyses, we trialled several purification schemes to remove the yolk, finding a magnetic nucleus immunolabelling and separation system (see methods) gave the purist yield of single nuclei. Using this nuclei extraction system, we processed 10-12 embryos for each sample in each of two separate experiments, determining mRNA levels and chromatin accessibility in individual nuclei by combined snRNA/ATACseq using a split-pool barcoding approach (Rosenberg et al., 2018, Zhu et al., 2021). After applying the RNA and ATAC quality-control criteria described in the Methods, we retained 24,282 nuclei with high-quality paired snRNA-seq and snATAC-seq profiles (Fig. S1B). The two modalities were integrated using Seurat/Signac Weighted Nearest Neighbor (WNN) analysis, followed by graph-based unsupervised clustering, which identified 10 distinct cell clusters (Fig. 1B). Based on RNA expression of established marker genes, 8 clusters were assigned to major embryonic cell types (Fig. 1C).

Using integrated snRNA/snATACseq, as expected at this stage of development we identified cell types from all the 3 main germ layers, ectoderm, mesoderm and endoderm, that closely resembled those of a Xenopus *tropicalis* cell atlas (Briggs et al., 2018). To assess the contribution of each modality, we also visualised the data using RNA-only and ATAC-only dimensional reductions. The RNA-only representation largely preserved the 10-cluster structure, whereas the ATAC-only representation showed broad separation of major cell identities but with fewer clearly resolved groups (Fig. S1C). This indicates that the ATAC modality captured biologically meaningful chromatin accessibility differences between cell populations, although with lower cluster resolution than the RNA modality. Superimposing developmental timepoints and treatment conditions onto the WNN UMAP showed the clear spatial separation of ASCL1-expressing cells from WT cells after 1.25hr and 2.5hr Dex treatment compared to WT GFP treated controls (Fig. 1B).

By comparing different cell clusters and timepoints in ASCL1-GR and WT GFP embryos, we revealed a widespread response to ASCL1 activation across multiple tissues that was confirmed by differentially expressed gene analysis (Fig. 1D,E). We found ASCL1 activation upregulated 899 genes distributed among all the 8 cell types at 1.25hr, increasing further to 1416 genes across all cell types at 2.5hr. As expected, the largest number of ASCL1-upregulated genes was found in neuroectoderm reaffirming the ability of ASCL1 to drive gene expression in neuroectoderm (Talikka et al., 2002, Hardwick and Philpott 2018, Ali et al 2014). However, changes in snRNA/ATACseq in response to ASCL1 activation were evident across all the major cell lineages, indicating major changes in gene expression and accessibility even in tissues not previously shown to respond to ASCL1 overexpression by upregulation of specific neural markers (Talikka et al., 2002, Hardwick and Philpott 2018, Ali et al 2014).

In addition to upregulated genes, we also found 1390 genes that were downregulated in response to ASCL1 activation across the 8 cell types at 1.25hr, decreasing to 515 genes by 2.5 hrs. Interestingly, mesoderm showed downregulation of 478 genes in response to ASCL1 activation, by far the greatest number downregulated genes in any cell type at 1.25hr. While this number of downregulated genes reduced to 57 at 2.5hr, the extent of gene downregulation was unexpected. We note that some target genes of ASCL1 that are induced in our study, such as Myt1 (later discussed in Fig. S3A), are well known transcriptional repressors of alternative non-neuronal gene programmes which help ASCL1 to drive neurogenesis (Bellefroid et al., 1996, Romm et al., 2005). However, the downregulation of large numbers of genes we observe after only 1.25 hours of ASCL1 activation indicates that indirect effects via induction of mRNA for subsequent protein expression of ASCL1-responsive downstream target repressors such as Myt1 are unlikely. Instead, our results, point to a potential direct repressive role of ASCL1 in this system. Together we conclude that ectopic ASCL1, in addition to inducing targets associated with neurogenesis in neuroectoderm, can also induce changes in mRNA expression of multiple genes across diverse tissue types in the early developing Xenopus embryo. Having established that all cell clusters show some response to ASCL1 activation, we next interrogated differences in those responses across cell clusters.

### ASCL1 induces neuronal and other gene programs across multiple tissues in all germ layers

Building on the known primary function of ASCL1 as a transcriptional activator (Lundie-Brown et al., 2025), we chose to focus specifically on all up-regulated genes to better understand the mechanism by which different cell types respond to ASCL1 by activating gene programmes. To understand if gene expression as a result of ASCL1 activation in different tissue types represents induction of primarily neuronal genes in all tissues, we undertook Gene Ontology (GO) enrichment analysis (Fig. 2A) on upregulated transcripts (Fig. 1E), comparing expression in different tissue clusters. Because functional annotation is more complete for human genes than for *Xenopus laevis*, *Xenopus* genes were mapped to their human orthologs before GO analysis, as described in the Methods.

**Figure 2.**
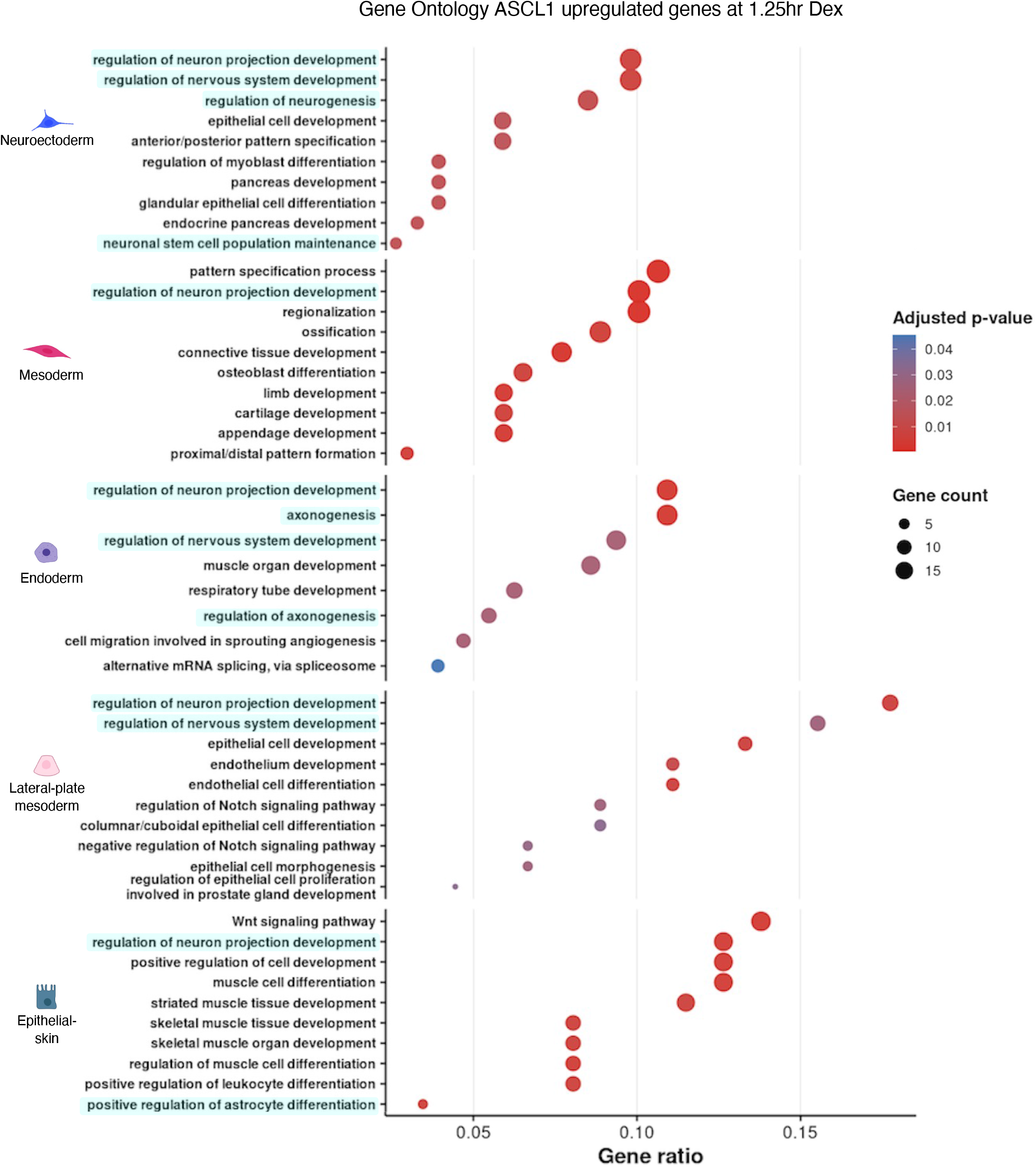
ASCL1 induces neuronal gene programs across many cell types. Gene ontology (GO) analysis of upregulated ASCL1 genes for each individual cell type after 1.25hr dexamethasone (Dex) treatment taken from Figure1E. Neuronal related gene programs are highlighted in blue.

In neuroectoderm, which is known to respond to ectopic ASCL1 by inducing neurogenesis (Talikka et al., 2002, Hardwick and Philpott 2018, Ali et al 2014), the most enriched GO terms for genes upregulated by ASCL1 activation were, as expected, related to neuronal programs (Fig. 2A). Surprisingly, neuronal-related terms were also enriched for genes upregulated in mesoderm, epithelial-skin, lateral-plate-mesoderm and endoderm cell types. This indicates that ASCL1 can induce genes related to neurogenesis in all three germ layers. However, GO analysis revealed that ASCL1 activated many additional non-neuronal gene programs, particularly in non-neuroectodermal tissues, for example several muscle related programs in epithelial-skin, and bone and cartilage terms in mesoderm (Fig. 2A). Our findings show that, although ASCL1 is transcriptionally active in all germ layers, it activates only minimally overlapping targets in different cell types in the developing embryo. This data suggests that the well documented pioneer activity of ASCL1 (Wapinski et al., 2013) does not result in clear convergent reprogramming to a neuronal identity in all embryonic tissue contexts.

### Asc1 activates overlapping and distinct target genes in different tissue cell types

Our gene expression and GO enrichment analysis suggests that all cell types respond to ASCL1 activation by upregulating large numbers of target genes, but neural beta tubulin-expressing neurons have previously been shown to be induced only in neuroectoderm in Xenopus embryos (Talikka et al., 2002, Ali et al 2014, Hardwick and Philpott 2018). To understand which neuroectodermally induced ASCL1 target genes were also being induced in the other cell types, we next investigated the activation of neuroectodermally-responsive ASCL1 targets in other cell types (Fig. 3A).

**Figure 3.**
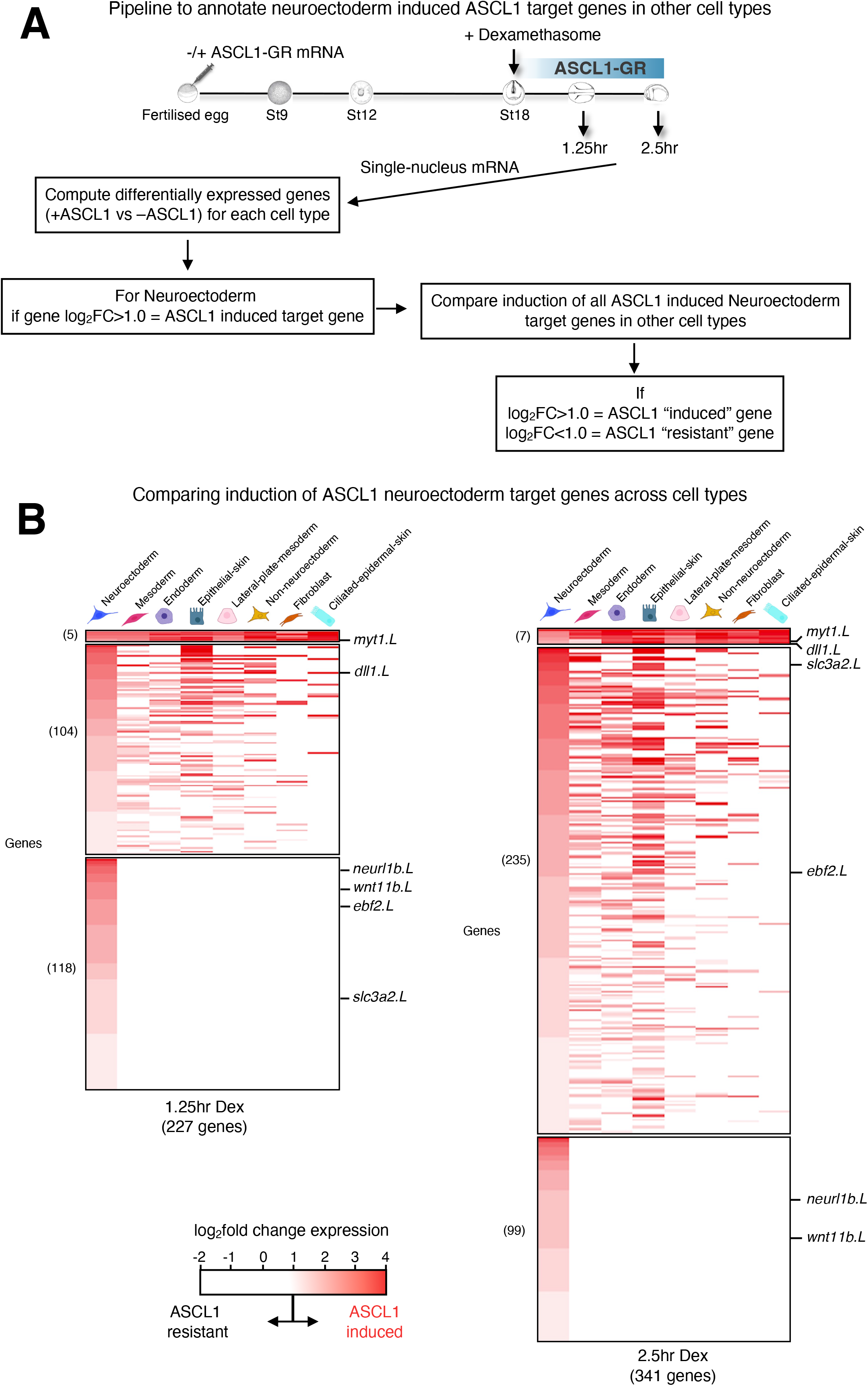
Comparing ASCL1 neuroectoderm target genes in other cell types identifies groups of differentially induced genes. A. Pipeline to identify ASCL1 induced and non induced (resistant) Neuroectoderm target genes in other cell types. B. Heatmap of log_2_ fold change in gene expression of ASCL1 induced Neuroectoderm target genes compared with induction in Mesoderm, Endoderm, Epithelial-skin, Lateral-plate-mesoderm, Non-neuroectoderm, Fibroblast and Ciliated-epidermal-skin at time points 1.25hr (left panel) and 2.5hr (right panel) after dexamethasone (Dex) treatment. Genes with a log_2_fold change >1.0 are annotated as “induced” by ASCL1 while those <1.0 are annotated “resistant” to being turned on by ASCL1.

**Figure S3.**
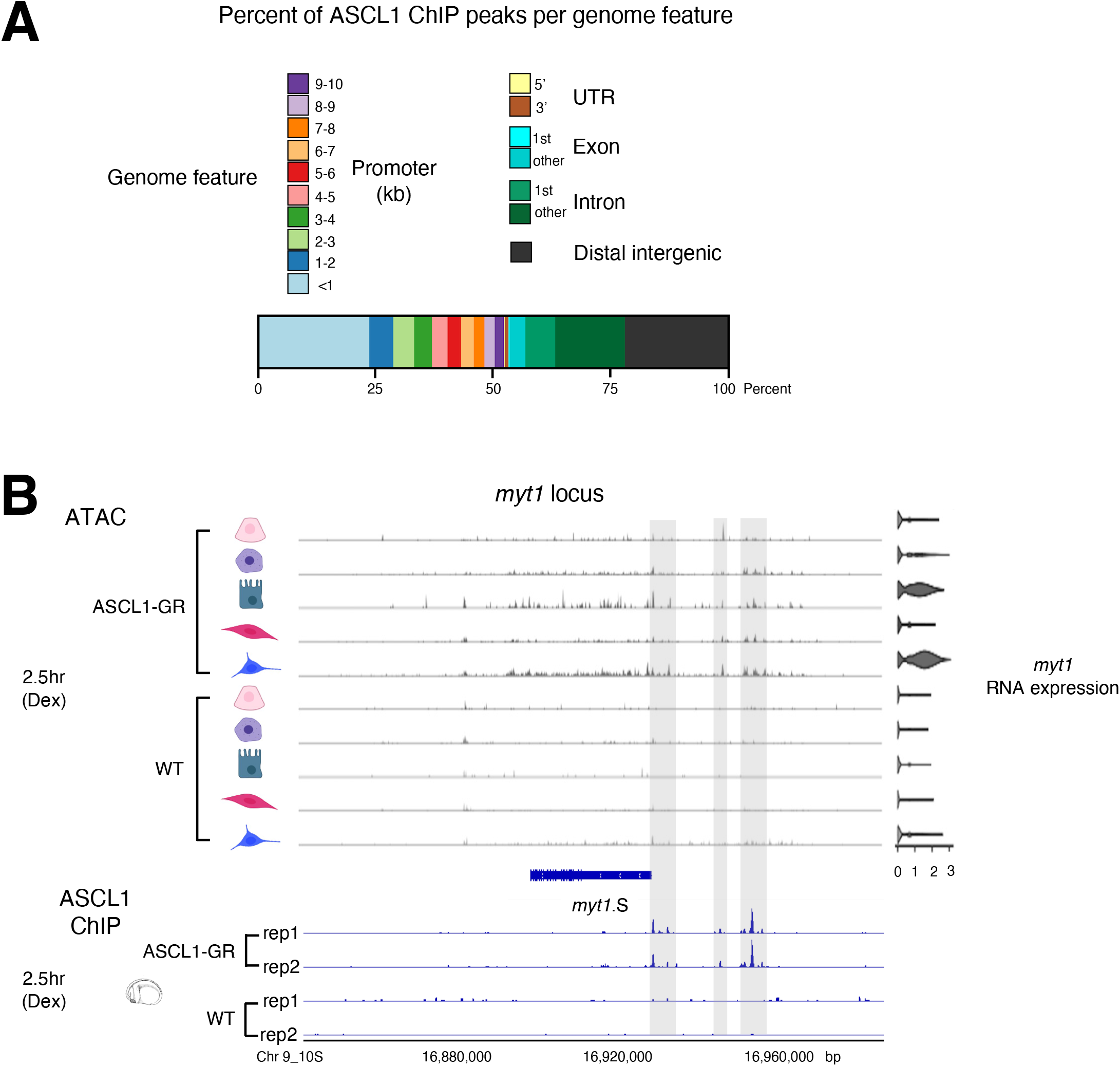
Ascl1 ChIP-seq data analysis. A. Plot of the percent distribution of 18,760 ASCL1 ChIP-seq peaks for each genome feature. B. Genome browser view of aggregated single-cell ATAC fragments and *myt1* RNA expression levels for 2.5hr dexamethasone (Dex) treated WT and ASCL1-GR samples from different cell types (top panel) and corresponding whole embryo ASCL1 ChIP-seq reads (bottom panel). Regions with ASCL1 ChIP-seq peaks are highlighted in grey.

Looking in more detail at our single nucleus mRNA analyses (Fig. 1, 2A), after 1.25hr of ASCL1-GR activation we identified a total of 227 ASCL1 upregulated genes that showed a log_2_fold-change greater than 1.0 in neuroectoderm compared to WT control. (Fig. 3B). We then undertook further analysis on this gene set activated by ASCL1 in neurectoderm. Comparing neuroectoderm-induced genes with induction of those same genes in mesoderm, epithelial-skin, endoderm, lateral-plate-mesoderm, non-neuroectoderm, ciliated-epidermal-skin and fibroblasts (Fig. 3B), at the 1.25hr timepoint, we found that only 5 neuroectodermally-induced genes were also induced across all other cell types. In addition, 104 neuroectodermally induced genes were found to also be upregulated in at least one, but not all, of these other cell types, while 118 genes were found to be induced in neuroectoderm only. Comparable analysis after 2.5hr activation identified 341 ASCL1 genes induced in neuroectoderm, with 7 of these also induced across all other cell types, 235 induced in at least one other cell type, and 99 induced in neuroectoderm only. This confirms that a small minority of ASCL1 targets are universally activated across tissue types, while larger numbers of genes are either upregulated in neuroectoderm alone or in neuroectoderm and only a subset of other tissues. We next investigated what could explain differences in response of specific genes to ASCL1 activation in different tissues.

### The extent of chromatin opening by ASCL1 correlates with tissue-specific target gene activation

To understand how changes in gene expression relate to ASCL1’s ability to access sites in embryonic chromatin we carried out whole embryo chromatin immunoprecipitation sequencing (ChIP-seq) to detect ASCL1 binding sites across embryonic tissues after 2.5hr dexamethasone treatment. After standard ChIP-seq processing, we identified 18,760 high-confidence ASCL1 binding sites. Analysis of the genome-wide distribution of ASCL1-bound sites revealed approximately 50% were located within 10kb of a gene promoter while approximately 20% were found within gene introns and distal inter-gene regions respectively (Fig. S3A).

Many cell lineage-defining transcription factors, including ASCL1, have been reported to regulate gene expression programs by acting as pioneer factors, opening chromatin at their binding sites (Iwafuchi-Doi and Zaret, 2014). To investigate whether ASCL1 opens otherwise potentially inaccessible chromatin in different cell types, we examined chromatin accessibility after 2.5hrs of ASCL1 activation proximal to the Myt1 gene locus, which we had identified as an ASCL1 gene target across all tissues (Fig. 3B and Fig. S3B). ASCL1-GR activated expression of Myt1 across all tissue types but to differing extents (Fig. 2B, right). We also observed greater chromatin accessibility at three regions located upstream from the Myt1 promoter identified as ASCL1 binding sites by ChIPseq after ASCL1 had been activated compared to wild-type cells (Fig. S2B, left). However, comparing across tissue types, the increase in chromatin accessibility appeared to correlate with the level of Myt1 gene upregulation by activated ASCL1; neuroectoderm and epithelial-skin show the greatest ASCL1-dependent increase in accessibility proximal to Myt1 and the greatest Myt1 gene activation, while mesoderm, endoderm and lateral-plate-mesoderm cells show a much lower chromatin opening by ASCL1 and a much lower gene activation response. We next investigated whether the changes in chromatin accessibility we saw in response to ASCL1 activation were responsible for the differential activation of neuroectoderm target genes by ASCL1 in the different tissues.

To investigate whether ASCL1 has a differing ability to open chromatin at its binding sites in different tissue contexts, we again focused on ASCL1 target genes activated by Ascl1 in neuroectoderm at both 1.25hr and 2.5hr timepoints and compared their expression in neuroectoderm and mesoderm. Ascl1 target genes were classified into two sets: i) shared targets; genes that are activated in both neuroectoderm and mesoderm and ii) unique targets; genes that are activated only neuroectoderm but not in mesoderm. For each target gene set, we defined candidate regulatory regions as the 50 kb region upstream of the transcription start site together with intronic regions of the corresponding ASCL1 target. These gene-associated regions were then intersected with the filtered whole-embryo ASCL1 ChIP-seq peak set to identify ASCL1-binding regions linked to shared and unique targets. Finally, we used the paired snATAC-seq data to quantify chromatin accessibility around ChIP-defined ASCL1 binding sites proximal to these genes in both neuroectoderm and mesoderm cells, using Signac region-level accessibility matrices and heatmaps centred on the ASCL1 peak regions (Fig. 4A).

**Figure 4.**
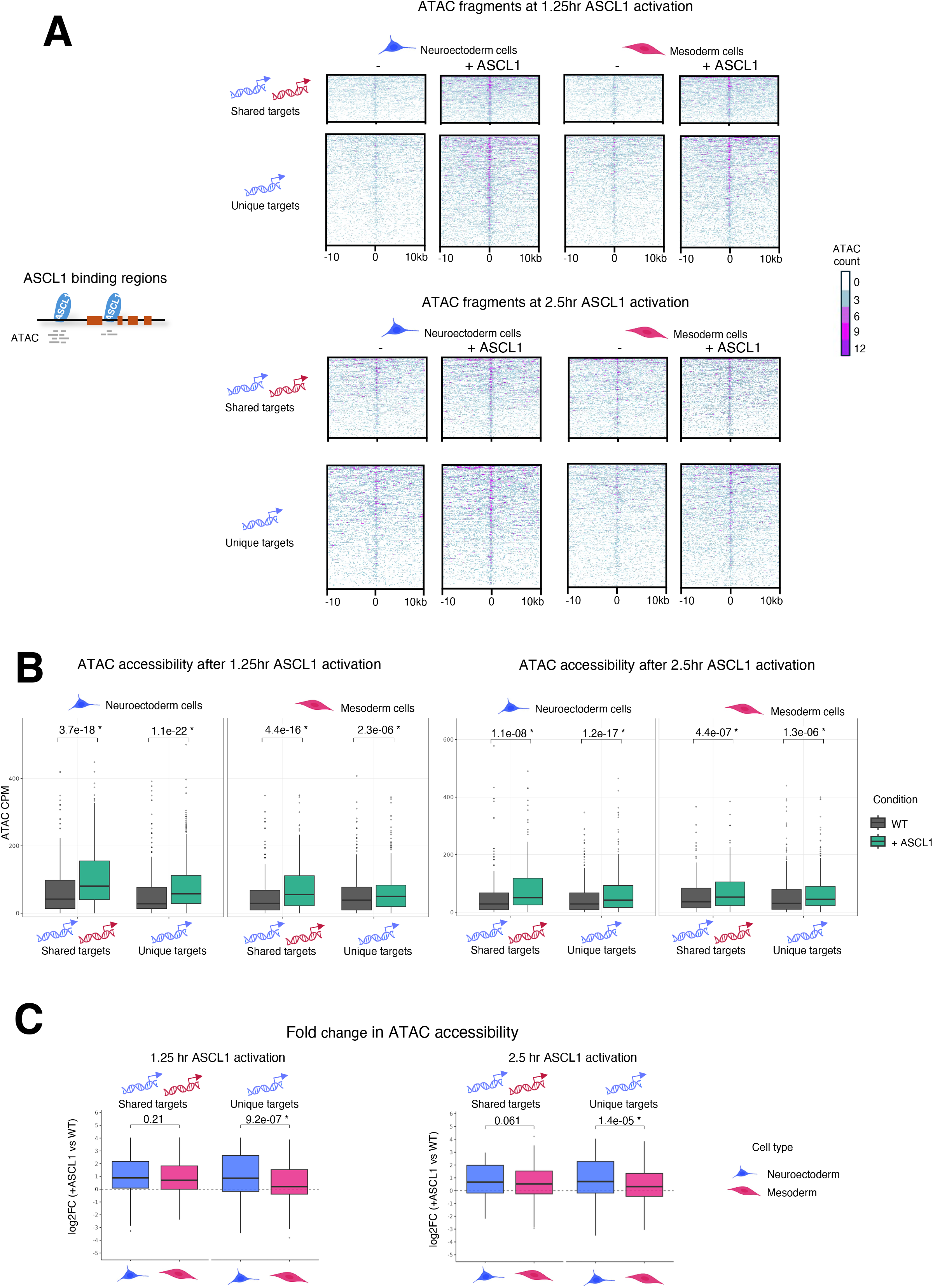
The amount of ASCL1-mediated chromatin opening correlates with tissue-specific target gene activation. A. Enrichment heatmap of single-cell ATAC fragments surrounding ASCL1 ChIP peaks from ASCL1 target gene activation groups induced in both Neuroectoderm and Mesoderm (shared targets) and Neuroectoderm only (unique targets), after 1.25hr and 2.5hr dexamethasone treatment. Each row depicts an ASCL1 binding site located up to 50kb upstream from the target gene transcriptional start site or an intronic region as shown in the diagram. While each panel are cells from Neuroectoderm or Mesoderm injected with ASCL1-GR mRNA (+ASCL1) or GFP mRNA (-) respectively. B. Box plot of the ATAC fragment counts per million (CPM) found across ASCL1 ChIP peaks (600bp window) for each of the sample panels described in (A) respectively. C. Plot of the log_2_fold change (FC) in ATAC fragment accessibility found at ASCL1 ChIP peaks of ASCL1 target gene activation group shown in (B) that are induced in both Neuroectoderm and Mesoderm (shared targets) and Neuroectoderm only (unique targets), after 1.25 and 2.5hr dexamethasone treatment for Neuroectoderm and Mesoderm cell types. Statistical *p-*values are from paired Wilcoxon sum test. * denotes significant *p-*value <0.05.

**Figure S4.**
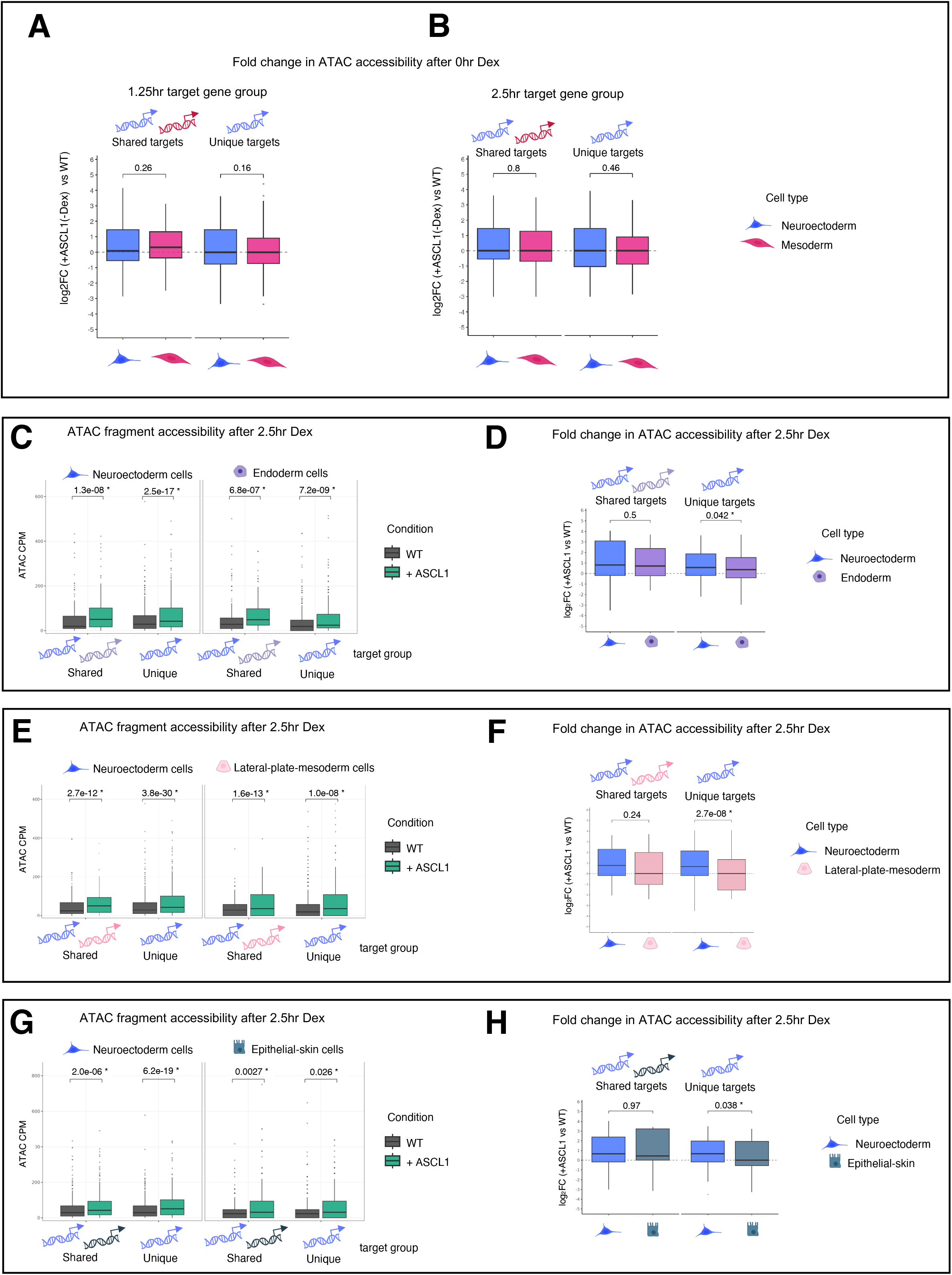
ASCL1-mediated chromatin opening correlates with tissue-specific target gene activation. A-B) Box plot of log_2_ fold change in ATAC fragment accessibility for Neuroectoderm compared to Mesoderm after 0hr dexamethasone treatment for ASCL1 ChIP peaks found at Neuroectoderm and Mesoderm target gene activation groups identified at 1.25hr (A) and 2.5hr (B) treatment. C-H) Box plot of the ATAC fragment counts per million (CPM) found across ASCL1 ChIP peaks for ASCL1 target gene activation groups induced in both tissues (shared targets) and Neuroectoderm only (unique targets), for cells from Neuroectoderm compared to Endoderm (C), Epithelial-skin (E) and Lateral-plate-mesoderm (G) after 2.5hr dexamethasome treatment. Corresponding log_2_ fold change plots in ATAC fragment accessibility found at the same ASCL1 ChIP peaks of ASCL1 target genes shown in (C, E and F) after 2.5hr dexamethasone treatment for Neuroectoderm compared to Endoderm (D), Epithelial-skin (F) and Lateral-plate-mesoderm (H). Statistical *p-*values are from paired Wilcoxon Sum Test. * denotes significant *p-*value <0.05.

We first examined whether pre-existing tissue-specific chromatin accessibility at ASCL1 binding sites of genes could account for unique or shared target gene expression. Without ASCL1 activation, we observed low but detectable enhancement in ATAC fragment density over background around ASCL1 binding sites associated with both shared and unique target gene sets in both tissues (Fig. 4A). Comparing chromatin at ASCL1-binding sites associated with shared and unique targets activatable by ASCL1, similar accessibility was seen at T=0 in ASCL1-GR injected embryos (no Dex) as was seen in GFP-injected control embryos (Fig S4A, B), demonstrating that the moderately accessible sites where ASCL1 favours binding are not generated by low level activation of ASCL1-GR without the addition of Dex. Instead, our data indicate that there is a more open configuration of ASCL1 binding sites compared to surrounding chromatin even prior to ASCL1 activation. We then investigated the effects of ASCL1 activation on chromatin accessibility.

As expected, ASCL1 activation in neuroectoderm cells significantly enhanced chromatin accessibility proximal to upregulated unique and shared target genes, (Fig. 4A neuroectoderm cell comparisons with quantification in Fig. 4B). Surprisingly, we saw also increased accessibility at ASCL1 binding sites in mesoderm cells in response to ASCL1 proximal to target genes regardless of whether activation of those genes was shared or unique to neuroectoderm (Fig. 4A, neuroectoderm/mesoderm cell comparisons with quantification in Fig. 4B). Thus, significant ASCL1-mediated enhancement of chromatin accessibility alone does not appear to be sufficient to drive gene expression of all neuroectodermal targets in mesoderm.

While opening of chromatin in both mesoderm and neuroectoderm was common to shared and unique targets, we next investigated the magnitude of changes to chromatin accessibility in both tissues in response to ASCL1. We calculated the fold change increase in ATAC fragments associated with ASCL1 binding sites near shared and unique targets in neuroectoderm and mesoderm cell clusters with and without ASCL1 activation (Fig. 4C). For shared target genes induced in both neuroectoderm and mesoderm, the increase in accessibility of associated ASCL1 binding sites was similar in neuroectoderm and mesoderm cells at both timepoints after ASCL1 activation. By contrast, ASCL1 targets unique to neuroectoderm showed a significantly higher increase in ATAC accessibility in neuroectoderm compared to accessibility in mesoderm after ASCL1 activation (Fig. 4C). Together, these data indicate that, while ASCL1 can enhance chromatin accessibility at neuroectoderm target genes with shared activation in both neuroectoderm and mesoderm, genes uniquely expressed in neuroectoderm is accompanied by greater ASCL1-dependent increase in accessibility compared to the more modest accessibility increase at sites in mesoderm associates with these unique genes.

To assess whether ASCL1 can open chromatin near neuroectodermal target genes in additional cell types, and whether this correlates with gene activation, we used a similar approach. The target genes activated by ASCL1 in neuroectoderm were now grouped according to whether they were also induced in endoderm, epithelial-skin, or lateral plate mesoderm (Fig. 3). We focused on these tissues because, like mesoderm, these clusters each contained at least 300 cells per timepoint, ensuring sufficient statistical power to detect differences in ATAC accessibility between conditions. We quantified ATAC-seq fragments at ASCL1 binding sites near neuroectoderm unique targets and targets whose expression was shared with the other tissues under analysis, comparing significant changes in accessibility near ASCL1 binding sites between conditions with and without ASCL1 activation across tissue pairs (Fig. S4C,E,G). Consistent with our observations in mesoderm, ASCL1 activation significantly increased chromatin accessibility around ASCL1 binding sites associated with unique and shared targets groups regardless of the cell type. We also evaluated whether the extent of increase in accessibility in the neuroectoderm compared to each of these other tissue types could explain the differences in gene activation (Fig. S4D,F,H), as we had observed in mesoderm. Again, consistent with our observations in mesodermal cells, after 2.5h activation of ASCL1, chromatin opening at unique targets was significantly greater in neuroectoderm compared to either endoderm, lateral-plate-mesoderm, or epithelial-skin.

Collectively, these results suggest that, while ASCL1 prefers to bind to moderately open sites and then can further open chromatin around both unique and shared targets in all tissues studied. However, the cell-type-specific activation of neuroectodermal target genes by ASCL1 is driven by the extent to which chromatin can be further opened by ASCL1 activation in a given tissue; in unique targets, the extent of further opening in non-neuroectodermal tissues appears to be insufficient to support gene activation.

### Chromatin accessibility identifies candidate transcription factors that may potentiate ASCL1 gene regulation

ASCL1 is capable of significant chromatin opening around its lineage-specific targets in neuroectoderm (Fig. 4 and Fig. S4). Given this, what mechanisms limit ASCL1 from successfully opening chromatin and activating these same neuroectoderm-only loci in alternate germ layers, such as mesoderm? Transcription factors rarely possess the structural means to initiate transcription or alter chromatin by themselves. Instead, they act within complex, combinatorial complexes of proteins requiring chromatin modifiers and partner co-factors to function (Reiter et al., 2017). We have observed that ASCL1 activation enhances chromatin accessibility some distance from its binding sites proximal to the *myt1* downstream target (Fig. 2B). We therefore reasoned that motif-specific co-factors could be needed to work with ASCL1 to activate neuroectoderm unique targets, while absence of these cofactors in mesoderm could result in ASCL1’s failure to activate the same genes in this context. One way to identify candidate cofactors required to activate ASCL1 targets in neuroectoderm is to look for transcription factor motifs that are enriched proximal to ASCL1-bound regulatory regions in this tissue. Differential accessibility of these ASCL1/co-factor motif-containing regions comparing across cell types could inform why ASCL1 activates some genes only in neuroectoderm, while other genes are activated in additional tissues such as mesoderm. We used this strategy to identify candidate cofactors required for gene expression in neuroectoderm (Fig. 5A).

**Figure 5.**
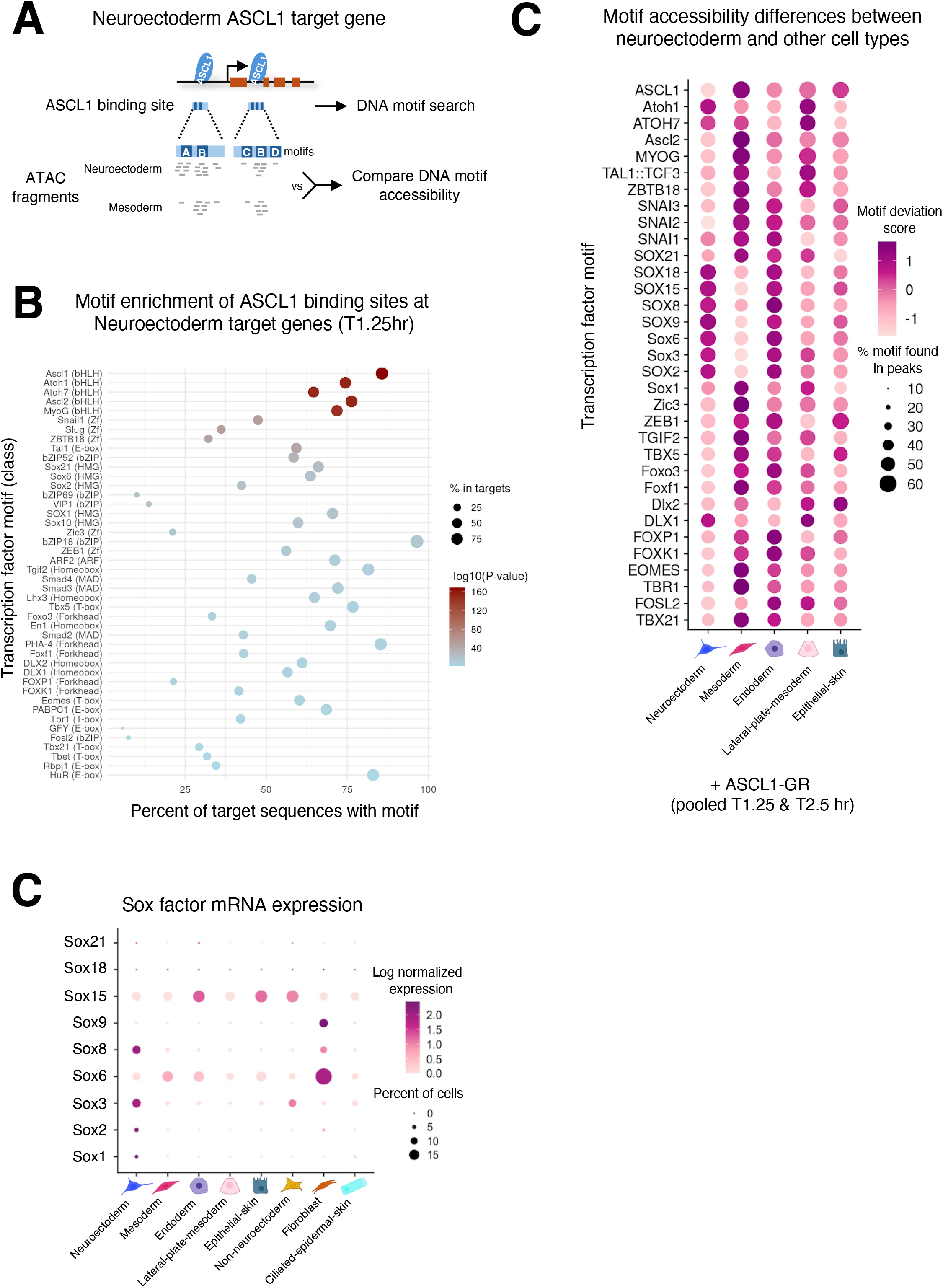
Motif accessibility analysis identifies potential signatures associated with ASCL1 gene regulation. A. Schematic of the approach used to identify transcription factor motifs enriched at ASCL1 target genes in Neuroectoderm and then compare the changes in ATAC accessibility of these motifs between Neuroectoderm and the other cell types. In this example motif A is more open at ASCL1 binding sites in neuroectoderm compared to mesoderm indicating motif A as a putative co-factor required by ASCL1 to turned on this gene in neuroectoderm which is lacking in mesoderm. B. Dotplot of the top 5 motifs from the top 10 motif classes found at ASCL1 binding sites of Neuroectoderm target genes after 1.25hr dexamethasone treatment. Homer was used to search 500 bp regions surrounding all Ascl binding sites located within 50kb region upstream from the transcriptional start site and intronic regions. For further details see Methods. C. ATAC accessibility dotplot of motifs from the top 10 classes identified in (B) between Mesoderm, Neuroectoderm, Endoderm, Lateral-plate-mesoderm and Epithelial-skin. ATAC motif deviation score of individual cells for each cell type was measured using ChromVar after pooling +ASCL1-GR mRNA (+ASCL1) cells from both 1.25 and 2.5hr dexamethasone treatments. A score>0 indicates more accessible, <0 less accessible than background. Only motifs with significant deviation scores >0 in at least one cell type have been plotted. D. Dotplot of log normalized single cell RNA expression for Sox factors in different cell types.

**Figure S5.**
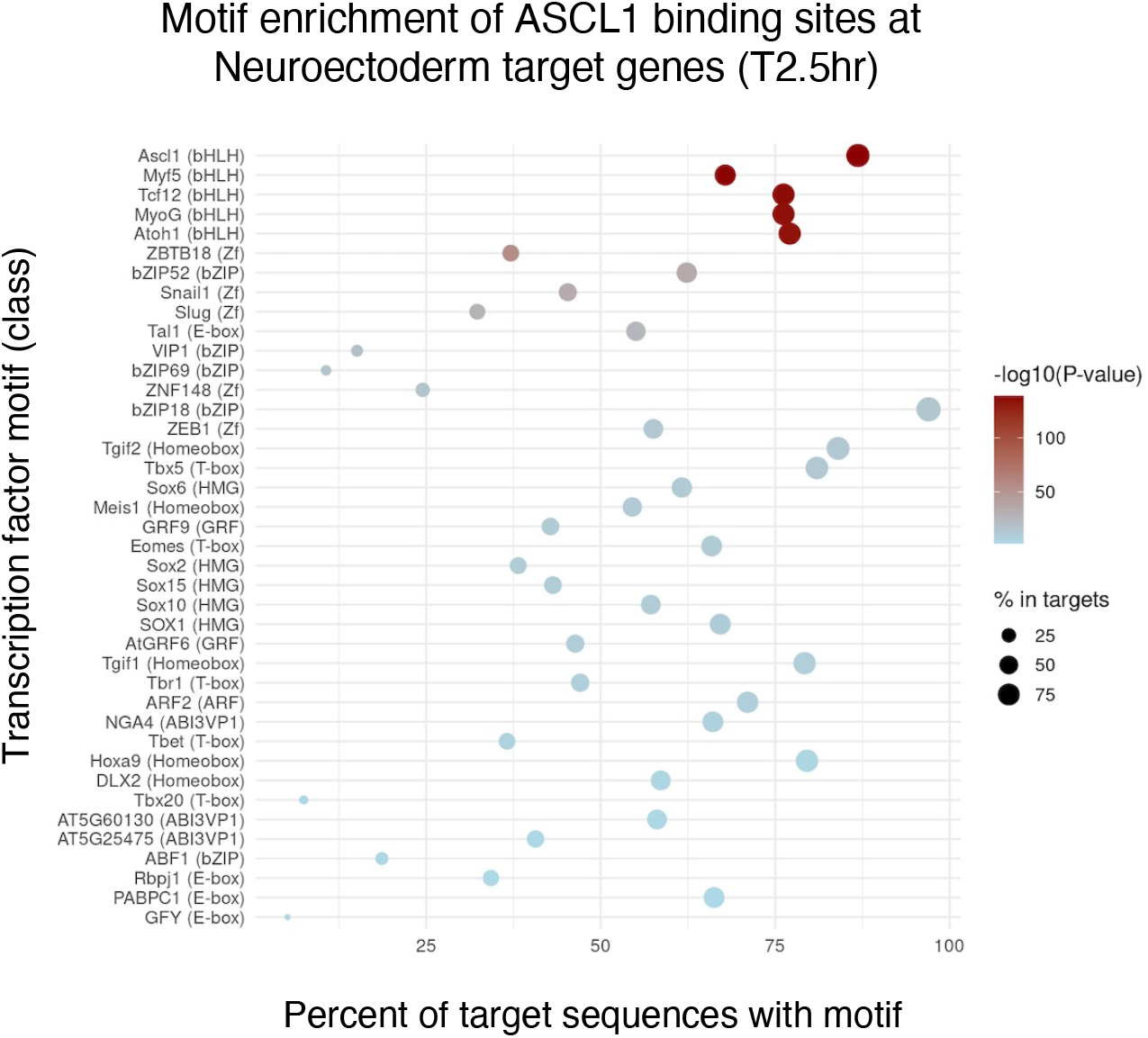
Motif analysis of ASCL1 binding sites surrounding neuroectoderm target genes. Dotplot of the top 5 motifs from the top 10 motif classes found at ASCL1 binding sites of Neuroectoderm target genes after 2.5hr dexamethasome treatment. Homer was used to search 500 bp regions surrounding all Ascl binding sites located within 50kb region upstream from the transcriptional start site and intronic regions.

We performed HOMER known motif enrichment analysis (Heinz et al., 2010) on ChIP-defined ASCL1 binding sites associated with neuroectoderm-induced genes after both 1.25hr and 2.5hr of ASCL1 activation (Fig. 5B and Fig. S5). Motifs associated with these regions were defined by intersecting high-confidence ASCL1 ChIP-seq peaks with proximal candidate regulatory regions assigned to genes activated in neuroectoderm, as described in the Methods. Broadly similar motif classes were enriched near ASCL1 binding sites proximal to neuroectodermally-activated genes at both timepoints. As expected, bHLH motifs were the most significantly enriched accessible sites found associated with identified ASCL1 binding sites of neuroectodermally activated genes; the ASCL1 motif was the most abundant bHLH factor motif found associated with over 80% of all sites. Other notable DNA elements found associated with these ASCL1 binding sites were Zinc-finger (Zf) motifs bound by the transcription factors Snail and Slug and High-Mobility-Group (HMG) box motifs bound by several SRY-box (Sox) factor proteins, in addition to proteins containing basic-leucine-zipper (bZIP) motifs.

Next, we identified which ASCL1-bound regulatory region motifs showed differential motif-associated accessibility after ASCL1 activation by comparing between different cell types, and then asked whether this differential motif accessibility correlates with a tissue-specific ability of the associated genes to be activated by ASCL1 (Fig. 5C). To ensure robust statistical confidence in identifying motif accessibility differences, for this analysis we pooled ASCL1 activated T1.25 and T2.5hr snATAC-seq data and only analysed cell clusters that had more than 700 cells. We then analysed the cell-by-peak accessibility matrix of these cell clusters, generated by quantifying the snATAC-seq fragment overlap with ChIP-defined ASCL1 peaks associated with all neuroectoderm activated genes. Motif identification within these peaks were identified using vertebrate transcription factor position frequency matrices from JASPAR (Ovek Baydar et al., 2026), and the chromVAR algorithm (Schep et al., 2017) was used to compute bias-corrected motif deviation scores for cells from each individual tissue. These deviation scores estimate whether regions containing a given motif are more or less accessible than expected in the tissue and conditions in which they are measured after correcting for technical biases such as GC content. Positive deviation scores indicate higher-than-expected accessibility of regions containing that motif, whereas negative scores indicate lower-than-expected accessibility (see Methods for further details).

Differential chromVAR analysis showed that, after ASCL1 activation, there was substantial cell-type-specific variation in motif-associated accessibility at the ASCL1-binding regions proximal to neuroectoderm activated genes. For example, we saw increased accessibility of several Sox-family motifs associated with ASCL1 sites at activated genes in neuroectoderm cells (Fig. 5C). These motifs associated with specific Sox factors are, however, not equally accessible in other tissues after ASCL1 activation. For instance, several Sox binding motifs associated with ASCL1 binding sites at activated genes in neuroectoderm are generally poorly accessible in mesoderm. Analysis of snRNAseq expression of these Sox factors in the different cell types also reveals strikingly different expression patterns; Sox factors such as Sox6 show higher expression in endoderm and mesoderm than in neuroectoderm, while Sox1, Sox2, Sox3 and Sox8 all show considerably higher expression in neuroectoderm than in mesoderm, endoderm, lateral-plate-mesoderm, or epithelia skin (Fig. 5D).

Taken together, our data show that there are over a hundred genes that are upregulated by ASCL1 in neuroectoderm that are not activated in mesoderm, and that Sox motifs proximal to ASCL1 binding sites near these genes are more highly accessible in neuroectoderm than in mesoderm. Moreover, several Sox factors are more highly expressed in neuroectoderm. Therefore, we hypothesised that one or more of these Sox factors are required for induction of ASCL1 target genes in neuroectoderm tissue and their absence from mesoderm results in failure to activate many neuronal targets of ASCL1 in this tissue.

### Co-expression of Sox3 with ASCL1 activates previously unresponsive ASCL1 target genes in mesoderm

Sox family proteins are a conserved group of transcriptional regulators defined by the presence of a highly conserved HMG DNA binding domain, first identified in Sry, a crucial factor involved in mammalian male sex determination (Gubbay et al., 1990, Sinclair et al., 1990). Vertebrate genomes contain approximately 20 Sox family members with several playing critical roles in the development of the neural crest and CNS in mammals and *Xenopus* (Schock and LaBonne, 2020, Rocha et al., 2014). Of the three Sox members we identified as potential ASCL1 co-factors in neuroectodermal gene expression (Fig. 5) whose motifs are less accessible and expression absent in mesoderm (Fig. 5D), we focussed on members of the SoxB1 family because Sox2 and 3 have known overlapping functional roles in the maintenance of neural precursor cells (NPC) (Graham et al., 2003) and early CNS formation in Xenopus, whereas Sox8 has been shown to be involved in the later specification of neural crest in *Xenopus* (O’Donnell et al., 2006). While little previous direct association with ASCL1 has been reported for these three Sox factors, we hypothesised that because ASCL1 drives neural precursor cells to differentiate, a SoxB1 family member with overlapping expression in NPCs is a good candidate ASCL1 co-factor.

To investigate a potential role for Sox3 in potentiating a neuronal response to ectopic ASCL1 in non-neuroectodermal tissues, we used *Xenopus* animal cap explants derived from blastula-stage embryos. Naïve animal caps are derived from ectoderm and develop into mucociliary epidermis when cultured to the equivalent stage 16 to 24 of embryo neurulation (Grunz and Tacke, 1989, Angerilli et al., 2018). Animal cap explants can be converted into mesoderm tissue by treating cultures with the potent mesoderm inducer Activin A (Smith et al., 1990). Thus, we have a convenient well-controlled system to investigate gene expression in response to ASCL1 in two different tissues (Fig. 6A).

**Figure 6.**
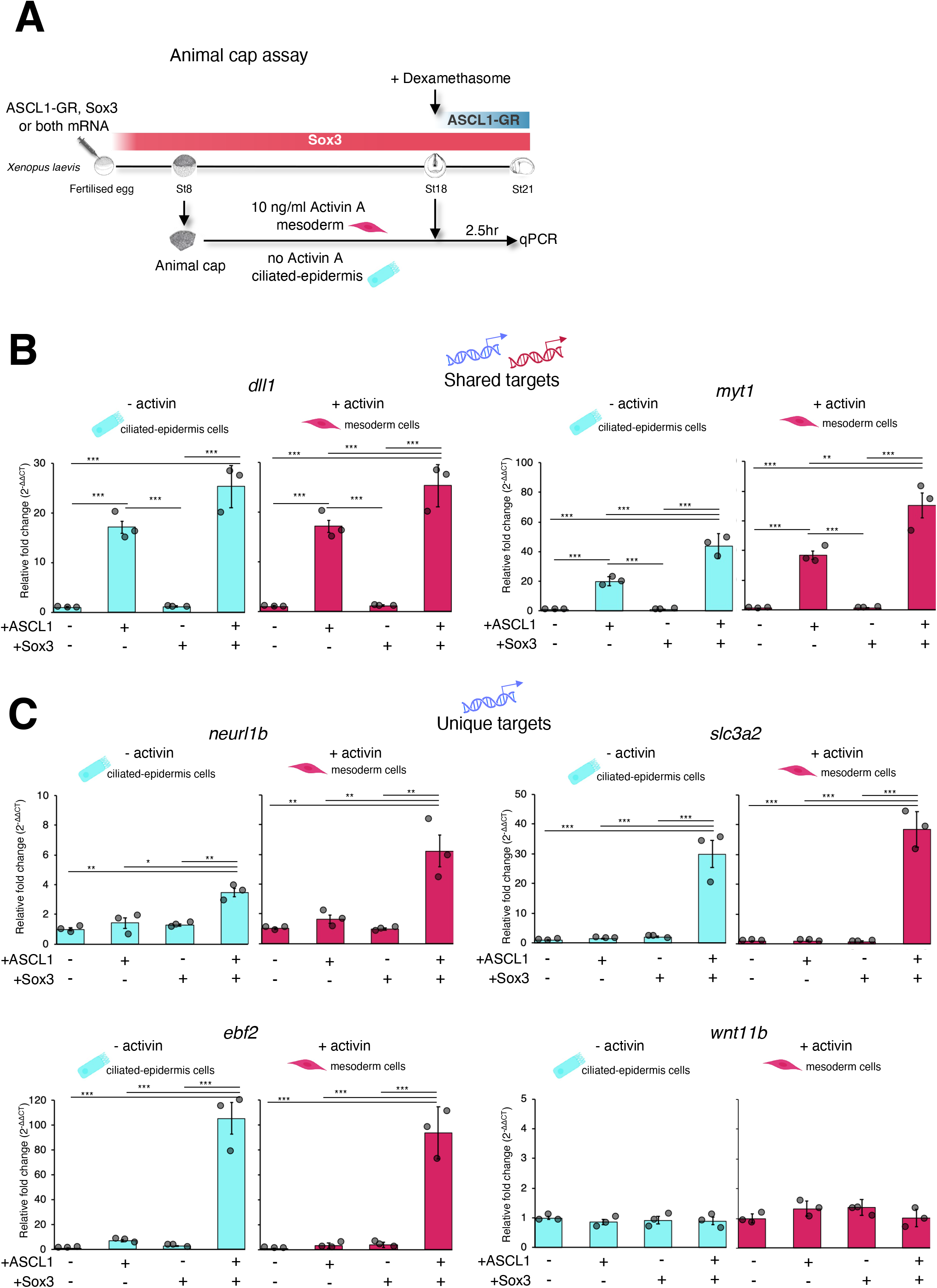
Sox3 rescues resistant ASCL1 target genes in Mesoderm. A. Schematic of the animal cap assay to test the co-expression of ASCL1 with Sox3 on gene expression changes in Mesoderm (+Activin A 10ng/ml) and Ciliated-epidermis (-activin) tissue. B & C. Gene expression fold change in mesoderm animal cap assay for different combinations of Sox3 and ASCL1 mRNA injections relative to non-injected control caps (WT). Panels in (C) are ASCL1 shared target genes induced in both Neuroectoderm and Mesoderm while (D) are unique target genes only induced in Neuroectoderm and not Mesoderm. Bars represent standard error mean of three biological replicates and statistical significance was determined by ANOVA with Tukey test *p<0.05, **p<0.01, ***p<0.001.

**Figure S6.**
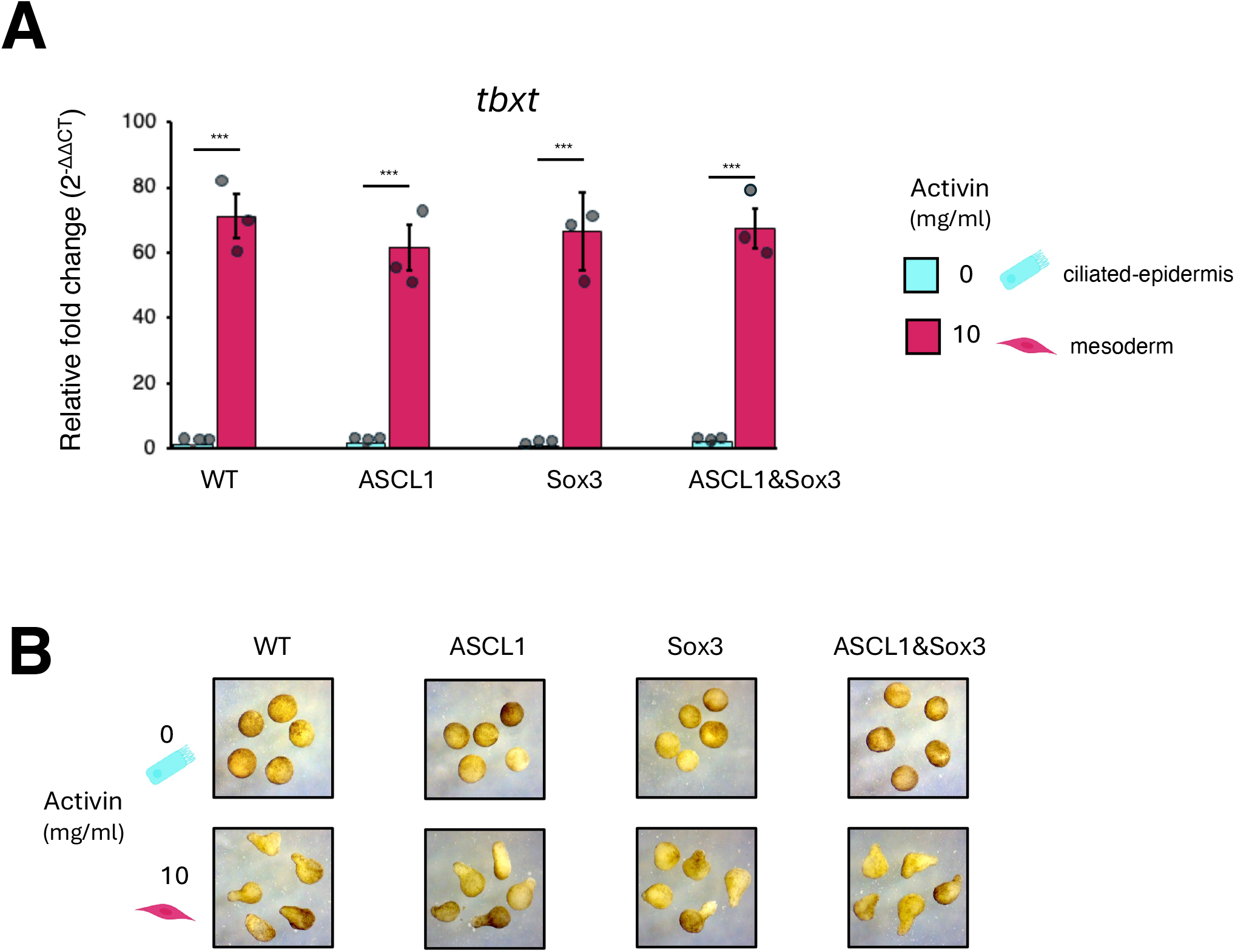
Quality control analysis for mesoderm animal cap assay. A) Gene expression fold change of the mesoderm marker gene *tbxt* with animal caps treated with (10mg/ml) and without (0) activin and injected Sox3 and ASCL1 mRNA as indicated. Gene expression is plotted relative to non-injected control caps (WT) without activin (0). Bars represent standard error mean of 3 biological replicates and statistical significance was determined with Wilcoxon Rank Sum Test *p<0.05, **p<0.01, ***p<0.001. B) Brightfield microscopy images of animal caps after treatment with (10mg/ml) and without (0 mg/ml) activin and injected Sox3 and ASCL1 mRNA as indicated. WT are non-injected control caps.

*Xenopus* embryos were microinjected with mRNA expressing either Sox3, ASCL1-GR or both together. At blastula stage, naive animal cap explants were isolated and cultured with or without 10 ng/ml Activin A. At a time equivalent to embryonic stage 18, animal caps were treated with dexamethasone to activate ASCL1-GR, and after 2.5 hr of Dex treatment, animal cap mRNA was isolated and gene expression was analysed by quantitative PCR (qPCR). Animal caps treated with Activin A significantly increased expression of the mesoderm marker *tbxt* (Fig. S6A) and showed a characteristic cap elongation (Smith et al., 1990), confirming their conversion into mesoderm tissue (Fig. S6B).

To assess if co-expression of Sox3 influences the expression of ASCL1 target genes in either a mesodermal or cillated epidermal tissue environment, we first measured the expression of *myt1* and *dll1*, genes that are both known to be activated by ASCL1 in neuroectoderm, mesoderm and ciliated-epidermal-skin in intact embryos (Fig. 3B). As predicted, overexpression of ASCL1 alone induced robust expression of both *myt1* and *dll1* in both mesodermal and ciliated-epidermal animal caps (Fig. 6B). Co-expression of ASCL1 with Sox3 resulted in further significant increases in RNA expression of both *myt1* and *dll1* genes in both tissues (Fig. 6B), showing that Sox3 can further enhance shared ASCL1 targets genes across tissues.

Next, we identified four ASCL1 target genes that are normally only expressed in neuroectoderm in response to ASCL1 (unique targets) (Fig.s 3), and that have adjacent Sox3 and ASCL1 motifs in their putative regulatory regions. RT-qPCR analysis of *neurl1b, slc3a2, ebf2* and *wnt11b* in animal caps in response to ASCL1 activation showed that ASCL1 alone fails to upregulate these unique targets in either mesodermal or ciliated-epidermal animal caps (Fig. 6C). Similarly, overexpression of Sox3 alone also fails to upregulate *neurl1b, slc3a2 ebf2 or wnt11b* in either animal cap tissue context. However, *neurl1b, slc3a2* and *ebf2*, were upregulated between 4 and 100-fold in mesoderm and ciliated-epidermal explants when ASCL1 and Sox3 are co-expressed, compared to uninjected controls. The *wnt11b* gene failed to be upregulated by either ASCL1, Sox3 or the combination. Taken together, this shows that expression of specific neuronal genes that are activated by ASCL1 only in neuroectoderm can be activated in other tissues by the addition of a Sox co-factor that has binding sites in close proximity to those of ASCL1, while other genes may have additional regulatory mechanisms to control tissue-specific expression.

## Discussion

We and others have previously shown that ectopic expression of ASCL1 in the developing *Xenopus* embryo can reprogramme dorsal ectoderm tissue to express neural markers such as β-III tubulin, but neural marker expression is not observed in mesoderm or endoderm tissue (Talikka et al., 2002, Ali et al 2014, Hardwick and Philpott 2018). This points to a difference across tissues in their competence to respond to an ectopic potent reprogramming transcription factor by promoting neurogenesis. To explore the nature of this difference in competence, in this study we investigate ectopic ASCL1 reprogramming activity across diverse tissues using single-nucleus multi-omics in *Xenopus* embryos during late stages of neurulation when the neural plate folds to form the neural tube.

Cells of the neuroectoderm lineage show the most extensive response to ASCL1 by activating the greatest number of genes after 1.25 and 2.5 hrs of activation compared to other cell lineages (Fig. 1E). As expected, Gene Ontology enrichment analysis of ASCL1-responsive genes reveals that neural pathways are among the top enriched terms in neuroectoderm (Fig. 2A). Despite their failure to express well established neural markers, Gene Ontology analysis shows that a proportion of the induced genes in all other cell types are also involved in neural pathways. This also could reflect engagement of a partial neural programme or the reuse of these same genes in other pathways. For example, neurons and pancreas share similar transcriptional networks and gene programs (Arntfield and van der Kooy, 2011), which may partly explain the activation we see of neural pathways in endoderm and of pancreatic pathways in neuroectoderm (Fig. 2A).

In addition to genes that may be associated with neurogensis, we also see the induction of gene programs associated with a variety of non-neuronal cell types in all tissue clusters, for instance muscle-related pathways in epithelial-skin (Fig. 2B). Muscle and cardiac off-target gene programs have been previously reported for ASCL1 reprogramming of mouse embryonic fibroblasts (Treutlein et al., 2016) and may result from promiscuous binding to sites more usually occupied by other bHLH factors such as MyoD, a core regulator of myogenesis (Davis et al., 1987). Indeed, the ability of ASCL1 to initiate changes in transcription of myogenic genes has been employed to enhance the efficiency of cardiac reprogramming (Wang et al., 2022).

Our results indicate that the varied gene expression responses and gene programs we see induced across the different tissues in response to ectopic ASCL1 likely reflect the cell intrinsic differences that constrain ASCL1 activity across different lineages, and also indicate that careful control of activated pathways is needed for heterologous cell types to adopt a neural identity. This cell intrinsic control on gene regulation in specific tissues even in the presence of a potent reprogramming factor is likely to underpin why previous studies have shown that ASCL1 induces neuronal marker expression only in neuroectoderm and not in mesoderm and endoderm tissue in the developing *Xenopus* embryo (Talikka et al., 2002, Ali et al 2014, Hardwick and Philpott 2018). Rather, the conflicting gene expression programs activated by ASCL1 in non-neuroectodermal tissues may confound re-enforcement of an imposed neuronal identity; instead, these ASCL1-activated genes associated with a variety of lineages cannot form an organised network to enforce any single new developmental trajectory This may explain why, superficially, mesoderm and endoderm tissue appear normal in embryos expressing ectopic ASCL1 despite ectopic ASCL1 resulting in major transcriptional changes across tissues revealed by snRNAseq (Talikka et al., 2002, Ali et al 2014, Hardwick and Philpott 2018).

Along with robust transcriptional upregulation of targets, the activation of ASCL1 also resulted in the downregulated expression of a large number of genes across all cell types (Fig. 1D). Indeed, at 1.25hr activation there were more downregulated targets than there were upregulated targets across all tissues. The rapid response to ASCL1 activation indicates some form of direct regulation by ASCL1, rather than an indirect mechanism requiring ASCL1-mediated induction of transcriptional repressors. A previous study in *Xenopus* has shown that ASCL1 can also act as a transcriptional repressor by physically interacting and recruiting histone deacetylase 1 to VegT target genes to repress mesendoderm formation (Gao et al., 2016). While we chose in this study to further explore gene activation as the most established function of ASCL1 (Guillemot et al., 1993; Lo et al., 1991), the role of ASCL1 in gene repression, which may or may not be dependent on ASCL1 binding motifs, could be explored in the future.

Although we show that genes that are upregulated in neuroectoderm in response to ASCL1 can also be activated in other tissues, the response of individual genes is variable and only a small subset of ASCL1 targets are activated in all cell types investigated. In fact, the majority of ASCL1 targets that are activated in neuroectoderm are either also activated in just one or two additional cell types, or show expression restricted to the neuroectodermal cluster alone (Fig. 3). ASCL1 has previously been characterised as a pioneer reprogramming transcription factor capable of opening closed chromatin at key neural genes in non-neuronal cell types, such as fibroblasts (Wapinski et al., 2013, Wapinski et al., 2017, Zhou et al., 2026). In cells that resist reprogramming, such as keratinocytes, ASCL1 appears unable to open chromatin at these key neuronal gene targets (Wapinski et al., 2013). We therefore hypothesised that the differences we see in gene activation between cell types in response to ASCL1 could be due to changes in the opening of closed chromatin in tissues that would not normally see proneural transcription factors such as ASCL1, like mesoderm.

Rather than opening closed chromatin, in neurula tissues at both unique and shared ASCL1 targets, we saw that ASCL1 prefers to bind to sites that are already partially accessible across all cell types (Fig. 4A)). Moreover, analysis of further chromatin opening at these sites revealed that ectopic ASCL1 could further open chromatin at binding sites associated with its neuroectodermal targets in all the cell types investigated, irrespective of whether that gene was activated by ASCL1 or not in a specific tissue (Fig. 4 and Fig. S4). However, we found that the extent of additional chromatin opening at neuroectodermal-only unique targets was significantly greater in neuroectoderm than mesoderm, while further opening was similar in shared targets activated in both neuroectoderm and mesoderm. Therefore in the embryo, while ectopic ASCL1 prefers to bind to open sites and can further increase accessibility of those sites across different tissues, the extent of that additional opening appears to contribute to the difference in the response of targets across different cell types. This subtle but important control mechanism is revealed by carefully controlled cross-tissue comparisons afforded by the *Xenopus* embryo system.

It is well known that the ability of ASCL1 to reprogram different mammalian cell types can be enhanced by co-expressing specific co-factors. In particular, the neuronal transcription factors BRN1 and MYT1L enhance the conversion of mesodermal fibroblasts and endodermal hapatocytes into neuronal cells (Marro et al., 2011, Vierbuchen et al., 2010). In addition, ASCL1 mediated conversion of astrocytes to neurons has been shown to require its downstream target NEUROD4 (Masserdotti et al., 2015). We have taken a novel approach to identify factors that can drive ASCL1-mediated activation of neuronal targets in tissues that are otherwise refractory to their expression. Our detailed analysis of accessible ASCL1 binding sites in neuroectoderm activated genes revealed cell-type-specific variations in the accessibility of associated co-factor motifs across tissues (Fig. 5B). In particular, motifs for several Sox factor proteins were found to be highly enriched in open chromatin associated with ASCL1 binding sites linked to neuroectodermally activated genes in neuroectoderm cells. By contrast, these motifs were largely absent from open chromatin proximal to ASCL1 binding sites in cell types such as mesoderm and epithelial skin that to not respond to ASCL1 by adopting a neural identity.

While ASCL1 and Sox factors have well documented roles in neurogenesis, they are not well described to act together to execute the neuronal programme. However, Sox proteins have been shown to interact with a variety of partner proteins including Sox21 with the bHLH factor Ngn2 (Whittington et al., 2015) and Sox2 with the chromatin remodeller ATRX (Bunina et al., 2020), and these interactions are important to direct neural progenitor cells to mature neurons. Here we show that co-expression of ASCL1 with Sox3 further enhances expression of several shared ASCL1 targets in both mesoderm and ciliated epidermal animal caps that are normally expressed across tissue types. In addition, the combination of ASCL1 and Sox3 allows several otherwise neuroectoderm unique targets to now be activated in mesoderm and ciliated epidermal skin (Fig. 6C). This demonstrates that co-factors that enhance reprogramming of otherwise refractory cell types can be identified by studying tissue-specific availability of co-located motifs in chromatin, and that a combination of two factors can significantly enhance upregulation of neuronal targets in mesoderm and skin.

ASCL1 has been well characterised as a reprogramming factor in mature mammalian cell culture systems. These studies have generally focussed on characterisation of the outcomes of reprogramming that are measured days or even weeks after transcription factor activation.

ASCL1’s activity in an embryonic reprogramming setting and in particular, in vivo, is much less well understood. One particular strength of our approach is that we can compare the earliest responses to a uniform ASCL1-mediated fate challenge across multiple tissues at the same developmental stage. Limited studies into the reprogramming events occurring in the first 12 to 48 hrs in mouse embryonic fibroblasts have shown that ASCL1 opens sites of closed chromatin at key neural genes at the onset of reprogramming (Wapinski et al 2013, Wapinski et al., 2017). Chromatin of the early mouse embryo has been reported to be more dynamic than that of more mature cells (Bošković et al., 2014). We observed that sites accessed by ASCL1 are already partially accessible in embryonic chromatin in different tissues, while further opening at these sites correlated with tissue-specific gene expression (Fig. 4). This could reflect the more flexible genome of the developing embryos that must respond rapidly to fate-determining factors over mature cells where stability of identity is likely to be more important.

## Supporting information

SingleCellMultiOmics_Xenopus_analysis

## Acknowledgements

The authors would like to thank the Genomics Facility at the Cambridge Institute for sequencing, the European Xenopus Research Centre Portsmouth for the Sox3 construct, the Cambridge University Aquatics facility for frog husbandry, Leon Peshkin, Anastasiia Ivanova and Elena Makarenkova for suggestions with nuclei separation methods, Rosalind Drummond for help with western blot and all members of the Philpott lab for helpful suggestions and discussions.

## Author contributions

Conceptualisation, D.L., T.S., J.J. and A.P.; Preliminary investigation, T.S., F.C. and J.J. Methodology and Data collection, D.L., T.S. and A.P.; Formal data analysis, visualisation and curation, S.K.; Writing – original draft, D.L.; Writing – review & editing, D.L., T.S., S.K., F.C., J.J., and A.P.; Supervision, J.J. and A.P.

## Data and resource availability

Custom code associated with analysis in this manuscript has been deposited on GitHub at https://github.com/Philpott-lab/SingleCellMultiOmics_Xenopus

## Methods

### Plasmids and mRNA synthesis

Full length mouse ASCL1 (mASCL1) fused to glucocorticoid receptor (GR) amino acid 512-777 ligand binding domain (Hollenberg et al., 1993), Green fluorescent protein (GFP) and *Xenopus* Sox3.L from the European Xenopus Research Centre were cloned into pCS2+ plasmid. Capped RNAs were synthesised in vitro from linearized plasmid template using the SP6 MEGAscript kit (Invitrogen).

### *Xenopus laevis*, ASCL1 overexpression and mesoderm animal cap assays

*Xenopus laevis* embryos were obtained and grown in 0.1x Modified Barth’s Saline (MBS) by standard methods and staged according to (Nieuwkoop and Faber, 2020). For overexpression experiments fertilised embryos were injected with 1 ng of ASCL1-GR or GFP and 200 pg Sox3 mRNA as indicated. For ASCL1 Sox3 co-expression a mix of 1 ng ASCL1-GR with 200 pg Sox3 was injected. For multiome and ChIP experiments embryonic stage 18 embryos were treated with 10 uM dexamethasome and grown at room temperature (23°C) for 0, 1.25 and 2.5 hrs. For mesoderm explant assays animal caps from stage 8 embryos were dissected and transferred to 0.7x MBS media containing 10 ng/ml recombinant rat Activin A (Cell Guidance Systems). At stage 18 caps were treated with 10 uM dexamethasome for 2.5 hrs. After treatment multiome and cap samples were snap frozen in a dry ice ethanol bath and stored at −80°C. For ChIP samples were fixed with 1% formaldehyde at room temperature for 25 minutes, washed 4 times in growth media, once in HEG solution (50 mM Hepes-KOH pH 7.5, 1 mM EDTA pH 8.0, 20% glycerol) followed by snap freezing on dry ice.

### Western blotting

Samples containing 3 embryos were thawed in 60 ul ice-cold E1 buffer (50 mM Hepes-KOH pH 7.5, 140 mM NaCl_2_, 1 mM EDTA, 10 % glycerol, 0.5% Igepal CA-630 (Sigma), 0.25% Triton-X-100 (Sigma), 1 mM DTT, 1x protease inhibitor cocktail (Roche cat.11873580001)) by gentle pipetting. Then centrifuge 1000g for 2 min at 4°C. Collect supernantant containing the cytoplasmic sample. Wash pellet with 1 ml ice-cold E1 buffer and again with 1 ml of ice-cold E2 buffer (10 mM Tris pH 8.0, 200 mm NaCl_2_, 1 mM EDTA, 0.5 mM EGTA, 1x protease inhibitor cocktail). To make the chromatin sample resuspend the pellet in 40 ul Emille buffer (500 mM Tris pH 6.8, 500 mm NaCl_2_, 1% NP-40 (Sigma), 1% B-mercaptoethanol (Sigma), 1x protease inhibitor cocktail). Add SDS PAGE buffer to both cytoplasmic and chromatin samples and heat for 5 min at 95°C. Spin and load onto 4-12% NuPAGE gel and then transfer to membrane. A monoclonal antibody was used to detect ASCL1 (BD Pharmingen cat no. 556604) and polyclonal rabbit antibody to detect Histone H4 (Cell Signalling cat no. 2592).

### Quantitative real-time PCR (qPCR)

RNA was purified using the Qiagen RNeasy mini kit and 1 ug of RNA was made into cDNA using the Qiagen QuantiTect reverse transcription kit. The cDNA was diluted to 150 ul with MQW and 2 ul was analysed by qPCR in an Applied Biosystems StepOnePlus machine using Promega GoTaq qPCR mix with 1 uM primer mix. Thermal cycling conditions: 95°C for 2 minutes, then 40 cycles of 95°C for 3 seconds, 60°C for 30 seconds. For primer sequences, see Table 1.

**Table 1.**
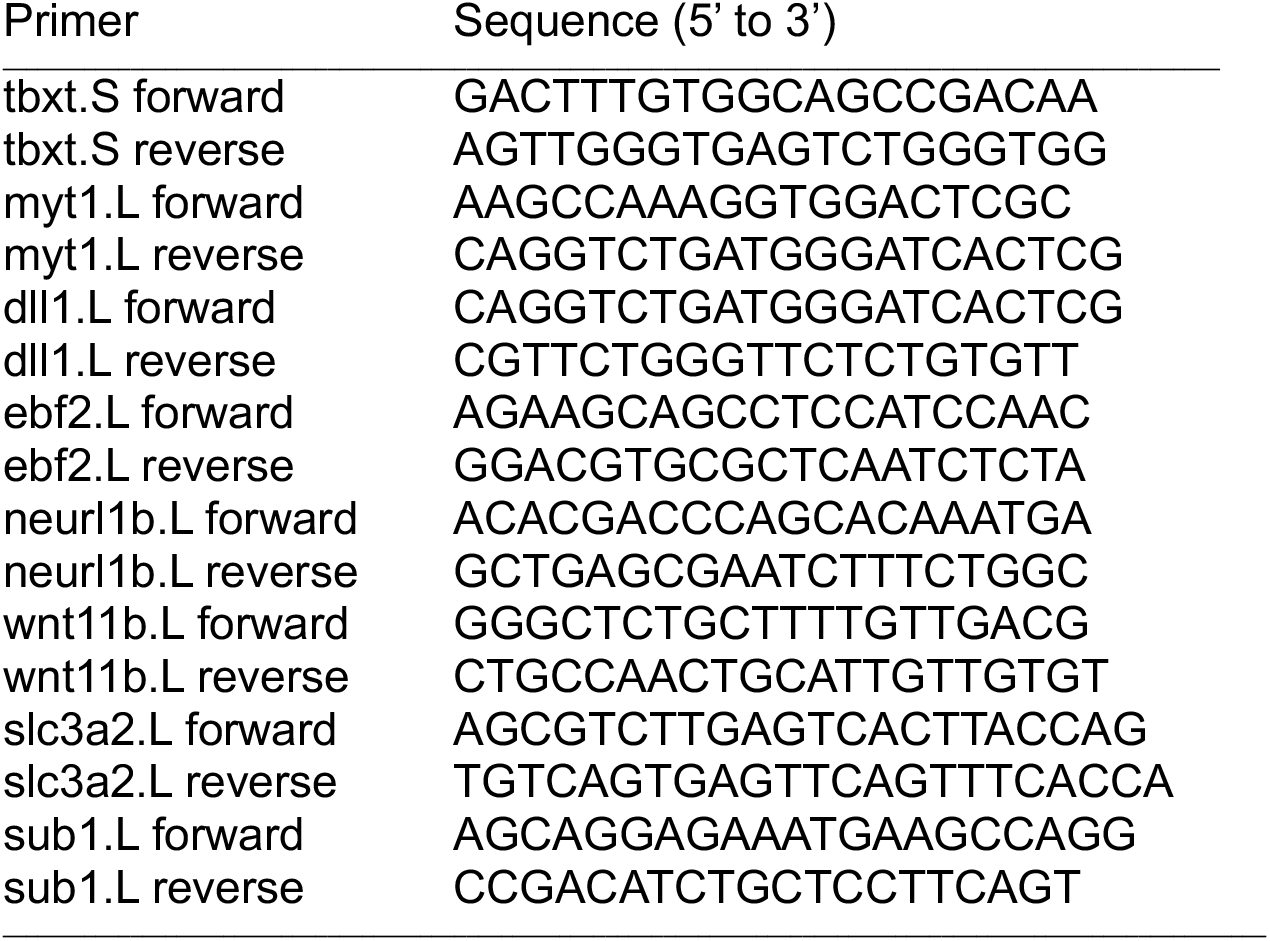
Primer sequences for qPCR.

### Chromatin Immunoprecipitation assay (ChIP)

Embryos were homogenized with buffer E1 (50mM Hepes-KOH pH 7.5, 140mM NaCl, 1mM EDTA pH 8.0, 10% glycerol, 0.5% Igepal CA-630, 0.25% Triton X-100, 1mM DTT, 1 x cOmplete Mini (Roche), 0.2mM PMSF). After a short spin (3500rpm, 2mins, 4°C), the pellet was resuspended in buffer E1 and kept on ice for 10 min. After a short spin, the pellet was resuspended in buffer E2 (10mM Tris pH 8.0, 200mM NaCl, 1mM EDTA pH 8.0, 0.5mM EGTA pH 8.0, 1 x cOmplete Mini, 0.2mM PMSF), and these steps were repeated three times. After a short spin, the pellet was resuspended in E3 (10mM Tris pH 8.0, 200mM NaCl, 1mM EDTA pH 8.0, 0.5mM EGTA pH 8.0, 0.1% Na-deoxycholate, 0.5% N-lauroylsarcosine, 1 x cOmplete Mini, 0.2mM PMSF). Sonication of chromatin was carried out on ice using Bioruptor (Diagenode), 4 cycles of high 5 mins (30s on/off). Centrifuge at 14K rpm at 4°C for 15 mins, then transfer the supernatant and add Triton X-100 to 1% final concentration.

Dynabeads M-280 (Dynal) 300ul were washed three times with 1ml PBS containing 0.1% BSA (PBS-BSA) and then 1,200ul PBS-BSA and 5ug anti-ASCL1 antibody (abcam, ab74065) were added and incubated 4°C overnight with rotation. Antibody-beads complexes were washed three times with 1ml PBS-BSA and once with 1ml buffer E2 supplemented with 1% TritonX-100. After the removal of E2 buffer, thawed sonicated chromatin and E3 buffer (to 5ml total volume) were added to the antibody-beads complexes and incubated at 4°C overnight with rotation. Then 5% of supernatant were removed as input fractions. Beads were washed 6 times for 5-10 minutes with pre-chilled RIPA buffer (50mM Hepes-KOH pH 7.5, 500mM LiCl, 1mM EDTA pH 8.0, 1% Igepal CA-630, 0.7% Na-deoxycholate, 1 x cOmplete Mini, 0.2mM PMSF), and twice with pre-chilled TEN buffer (10mM Tris pH 8.0, 1mM EDTA pH 8.0, 150mM NaCl, 1 x cOmplete Mini, 0.2mM PMSF). Beads were resuspended with 500 ul of STOP solution (40mM Tris pH 8.0, 10mM EDTA pH 8.0, 1% SDS) with 7.5ul Proteinase K (NEB, P8107S) and 25ul of 5M NaCl and were incubated overnight at 65°C.

After Phenol:Chloroform extraction and following ethanol precipitation, indexed libraries were generated using NEXTFLEX ChIP-Seq Kit (PerkinElmer) according to the manufacturer’s protocol. Libraries were sequenced using NovaSeq 6000, paired-end 50 cycles.

### ChIP data processing

Paired-end ChIP-seq reads were first assessed using FastQC v0.12.1, followed by adapter and quality trimming with fastp v0.23.4 (Chen et al., 2018) using paired-end adapter detection, poly-G trimming, a minimum base quality threshold of 20 and a minimum read length of 20 bp. Trimmed reads were aligned to the Xenopus laevis NCBI RefSeq genome assembly GCF_017654675.1 using Bowtie2 v2.3.5.1 (Langmead and Salzberg, 2012). Alignments were performed in paired-end mode using the parameters -N 0, --dovetail, --fr, --no-mixed, --no-unal and a maximum fragment length of 1000 bp. SAM files were converted to BAM format using SAMtools v1.9 (Danecek et al., 2021), and alignment statistics were assessed using samtools flagstat.

BAM files were filtered using Sambamba v0.6.8 to retain properly paired, mapped, non-duplicate reads without secondary alignment tags. Filtered BAM files were then sorted and indexed using SAMtools v1.9. Paired-end fragment size distributions were assessed using deepTools bamPEFragmentSize v3.5.1. CPM-normalised bigWig files were generated using deepTools bamCoverage v3.5.1 with a bin size of 10 bp, read extension enabled and an effective genome size of 2,864,785,220 bp.

Peaks were called using MACS2 v2.2.9.1 (Zhang et al., 2008) in paired-end mode (-f BAMPE) with the *Xenopus laevis* effective genome size set to 2.74079e+09 and --keep-dup all. For each ASCL1 ChIP-seq replicate, peaks were first called against the corresponding input sample. In addition, ASCL1-treated replicates were compared against control ChIP samples to identify and remove non-specific or background-enriched regions. To define a reproducible ASCL1 binding set, MACS2 peak calls were imported into DiffBind v3.16.0 (Ross-Innes et al., 2012), and a consensus peak set was generated by retaining peaks detected in at least two of the three ASCL1 ChIP-seq biological replicates. Peaks overlapping control-enriched regions were removed, and the remaining consensus peaks were further filtered by ChIP-seq signal, retaining only peaks with an average read count of at least 100 across ASCL1 replicates. This filtering resulted in a final set of 18,760 high-confidence ASCL1 binding sites. This final peak set was used for downstream genomic annotation, assignment to candidate target genes and integration with the single-nucleus ATAC-seq data.

### Single-nucleus RNA & ATAC multiome library preparation and sequencing

To prepare nuclei for single-nucleus multiome we used the magnetic nuclei isolation system from Miltenyi Biotec. Samples containing 10-12 embryos were thawed and dissociated on ice by resuspending in 1ml of ice-cold nuclei extraction buffer (Miltenyi Biotec) supplemented with 1x protease inhibitor cocktail (Roche cat.11873580001), 0.5U/ul ea Superase RNAse Inhibitor and RNAaseOUT. Once dissociated cells were centrifuged at 600g for 5 min at 4°C. Supernatant was removed and the greyish brown pellet was resuspended in 1 ml of separation buffer (1 vol nuclei extraction buffer, 6 vol 1x PBS (Gibco) supplemented with 200 ug/ml BSA (Invitrogen), 1x protease inhibitor cocktail, 0.5U/ul ea Superase RNAse Inhibitor and RNAaseOUT). Nuclei extract solution was then filtered through a 30-micron filter (pluriSelect) and centrifuged at 600g for 5 min at 4°C. Again, gently remove the supernatant and resuspend pellet in 450 ul separation buffer. Add 50 ul of anti-nucleus microbeads (Miltenyi Biotec), gently mix and incubate for 15 min at 4°C. After incubation add the bead sample to a pre-equilibrated LS column (Miltenyi Biotec) attached to the QuadroMACS magnetic stand. After the sample has run through wash the column twice with 1 ml of separation buffer. To elute the purified nuclei, remove column from the magnetic stand and collect nuclei by adding 1.0 ml fix buffer (1x PBS, 0.2% formaldehyde (Pierce), 1x protease inhibitor cocktail, 0.5U/ul ea Superase RNAse Inhibitor and RNAaseOUT). After 5 min incubation on ice fixation was stopped by adding 25 ul 1M Tris-HCl pH7.6 containing 0.1mg/ml BSA. Centrifuge 600g for 5 min at 4° C. Wash pellet once with 200 ul PBS solution (1x PBS containing 1x protease inhibitor cocktail and 0.5U/ul ea Superase RNAse Inhibitor and RNAaseOUT). Resuspend nuclei in PBS solution.

For ATAC transposase reaction 10 ul of each sample containing 75,000 cells were aliquoted in 0.2 ml PCR tubes containing 5 ul (25 uM) of loaded Tn5 transposase (Diagenode) with appropriate sample barcode as described in Zhu et al., 2021. Gently mix and incubated on ice for 5 min. Then add 35 ul of Tn5 transposase buffer (20 mM Tris-HCl pH 7.6, 10 mM MgCl_2_, 20% Dimethyl formamide, 0.1% Tween-20, 0.01% digitonin, 0.5U/ul ea Superase RNAse Inhibitor and RNAaseOUT). Gently mix and incubate samples for 30 min at 37°C. After incubation centrifuge 800g for 5 min at 4°C. Wash pellet once with 200 ul PBS solution. Resuspend final pellet in 16 ul PBS solution.

To each tube add 24 ul reverse transcription mix (1x Maxima H buffer, 0.5 mM dNTPs, 0.1mg/ml BSA, 400U Maxima Reverse H Minus Reverse Transcriptase (Thermo Scientific) and 0.5 U/ul ea Superase RNAse Inhibitor and RNAaseOUT) containing 2.5 µM ATAC matched barcoded primers as described in Rosenberg et al., 2018. The reverse transcription was then performed in a thermocycler with the following program: Step 1: 50°C × 10 min; Step 2 (3 cycles): 8°C × 12 s, 15°C × 45 s, 20°C × 45 s, 30°C × 30 s, 42°C × 2 min, 50°C × 5 min; Step 3: 50°C × 10 min. After the reaction, the nuclei were transferred and pooled on ice into a 1.5-ml tube prewashed with 5% BSA in 1x PBS. Then Triton-X-100 was added to a final percent of 0.075 and nuclei were pelleted at 800g for 5 min at 4°C.

For ligation-based combinatorial barcoding the pooled nuclei were resuspended in 1 ml of 1x NEBuffer3.1 (NEB) and transferred to 2,910 ul of ligation buffer (1x T4 ligase buffer (NEB), 0.35mg/ml BSA, 25 mM NaCl_2_, 40,000U T4 ligase (NEB), and 0.25 U/ul ea Superase RNAse Inhibitor and RNAaseOUT). Using a multichannel pipette, 40 ul of the ligation mix was distributed to the 96 wells of the first R02 barcode plate (Rosenberg et al 2018). The plate was sealed and placed on a Thermomixer (Eppendorf) and mixed for 30 sec at 37°C at 1200rpm. Then incubated for a further 30 min at 37°C at 300rpm. After incubation the plate was removed and 10 ul of R02 blocking solution (26 µM R02 blocking oligo 5’ ATCCACGTGCTTGAGAGGCCAGAGCATTCG in 1x T4 ligase buffer) was added to each well. The plate was re-sealed and mixed again for 30 sec at 1200rpm before incubating for an additional 30 min at 37°C. The nuclei from each well were then pooled into 1.5 ml tubes and pelleted at 800g for 5 min at 4°C. The second round of ligation was then carried out using R03 biotin tagged barcode plate (Rosenberg et al 2018) similarly to the first round, except that after the 30 min ligation reaction, blocking solution (26 µM quencher oligo 5’ GTGGCCGATGTTTCGGTGCGAACTCAGACC in 0.25 M EDTA) was added to quench the reaction. After 2 rounds of ligation-based barcoding, the pooled nuclei were resuspended in 1x PBS with 0.5U/ul ea Superase RNAse Inhibitor and RNAaseOUT, counted and aliquoted into 10 µl sublibraries containing 3,000 nuclei.

For each sublibrary nuclei were lysed and cross-links reversed by adding 10 ul of lysis buffer (10 mM Tris-pH 7.5, 1M NaCl_2_, 0.5 mM EDTA, 2% SDS) and incubating at 65°C for 5 hrs. Biotin barcoded cDNA molecules were purified by adding 80 µl of dynabead solution (10ul MyOne Streptavidin C1 dynabeads (Invitrogen) in 10 mM Tris-pH 7.5, 1M NaCl_2_, 0.5 mM EDTA) and incubating for 1 hr at 24°C at 1000rpm. Bound cDNAs were washed twice with 200 ul BW buffer (10 mM Tris-pH 7.5, 1M NaCl_2_, 0.5 mM EDTA) followed by 2 additional washes with 200 µl Tris-T solution (10 mM Tris-HCL pH7.5, 0.1% Tween 20). For incorporating a 5’ adaptor onto the streptavidin bound cDNA a 50 µl template switching mix (1x Maxima H buffer, 4% Ficoll, 1mM dNTPs, 2.5 µM TS oligo (AAGCAGTGGTATCAACGCAGAGTGAATrGrG+G where rG is RNA base), 500U Maxima H minus RT, 15U Superase RNAse Inhibitor and 20U RNAaseOUT) was added to each sublibrary. The bead solution was then incubated at 25°C for 30 minutes and then at 42°C for 90 minutes with mixing at 1000rpm. After the incubation beads were washed twice with 200 µl Tris-T solution.

Washed beads were resuspended in 50 µl of 1^st^ round PCR mix (1x Q5 reaction buffer, 0.1 mM dNTPs, 0.5 uM each RNA (AAGCAGTGGTATCAACGCAGAGT), Forward (CAGACGTGTGCTCTTCCGATCT) and ATAC (AATGATACGGCGACCACCGAGATCTACACTAGATCGCTCGTCGGCAGCGTC) primer, 1U Q5 High-Fidelity DNA polymerase (NEB)) and amplified with the following program: Step 1: 72°C × 1 min; Step 2: 95°C × 1 min; Step 3 (6 cycles): 98°C × 30 s, 67°C × 15 s, 72°C × 4 min; Step 3: 72°C × 1 min. PCR reactions were purified using a 0.85X ratio of SPRI beads (Beckman) and eluted sample was split equally into two PCR tubes. One PCR tube was used to amplify the RNA library while the other tube the ATAC library.

For the RNA library we determine the optimal PCR cycle to give 1/3^rd^ of a saturated signal by repeating the PCR using the RNA and Forward oligo, 1/10^th^ of the purified PCR reaction and EvaGreen (Biotium) in a qPCR machine with the following program: Step 1: 95°C × 1 min; Step 3 (30 cycles): 98°C × 30 s, 70°C × 15 s, 72°C × 3 min; Step 3: 72°C × 1 min. Using the optimal cycle we PCR amplified the remaining cDNA and purified the reactions using 0.85X ratio of SPRI beads. To normalize the amplified cDNA fragment lengths for sequencing we trimmed the libraries using Tn5 transposase (Diagenode) loaded with (5’-[Phos] CTGTCTCTTATACACATCT, 5’-TCGTCGGCAGCGTCAGATGTGTATAAGAGACAG) to a size range of 200-1000bp. Each trimmed sublibrary was then PCR amplified using a unique forward P7Trueseq index barcode (5’ CAAGCAGAAGACGGCATACGAGATXXXXXXGTGACTGGAGTTCAGACGTGTGCTCTTCC GATC) and reverse oligo (AATGATACGGCGACCACCGAGATCTACACTAGATCGCTCGTCGGCAGCGTCAG). Like the RNA libraries we also determine the optimal PCR cycle for the ATAC libraries using the ATAC and Forward oligos with a PCR 72°C extension time of 30 sec. Using the optimal cycle we PCR amplified the remaining ATAC sample using a unique forward P7Trueseq index barcode and reverse oligo. Purified RNA and ATAC libraries were than sequenced on a NovaSeq 6000 using 100bp paired end reads and a read depth of 25-30,000 reads per cell.

### Single-nucleus multiome data preprocessing

Raw split-pool single-nucleus RNA/ATAC sequencing data were processed using the Paired-Tag pipeline described by Zhu et al., 2021 (https://github.com/cxzhu/Paired-Tag). Briefly, cellular barcodes were extracted from read 2, while read 1 was used to assign RNA and ATAC reads after alignment to the *Xenopus laevis* reference genome v10.1. Demultiplexed reads from individual sublibraries were merged to generate cell-barcode gene-count matrices for the RNA modality and chromatin accessibility files for the ATAC modality. Because the original Paired-Tag workflow was developed primarily for well-annotated reference genomes, custom annotation formatting and additional in-house processing were used to generate *Xenopus laevis*-compatible RNA count matrices and ATAC fragment files for downstream Seurat/Signac analysis.

### Gene-structure-window feature definition for scATAC-seq analysis

For the ATAC modality, we evaluated alternative feature definitions because peak-based feature construction from shallow split-pool scATAC-seq data in Xenopus laevis produced sparse and less stable matrices. We compared standard peak-based features, fixed-width genomic bins and gene-structure-window features. Gene-structure-window features were defined using annotated gene bodies together with upstream regulatory windows, generating an interpretable feature space linked to gene structure. Feature definitions were benchmarked by comparing ATAC-derived embeddings and clusters against RNA-derived cell-type annotations from the paired multiome data. Gene-structure-window features showed improved agreement with RNA-defined cell identities compared with peak-based features and were therefore selected for the final ATAC feature matrix for downstream dimensionality reduction and multiome integration.

### Single-nucleus multiome data quality control and multiome integration

Single-nucleus multiome analysis was performed in R using Seurat v5 (Hao et al., 2023) and Signac v1 (Stuart et al., 2021) RNA and ATAC modalities were quality-controlled independently, and only cells passing filters for both modalities were retained. RNA data were normalised using SCTransform and reduced by principal component analysis. ATAC data were normalised using TF-IDF transformation and reduced by latent semantic indexing. The RNA and ATAC modalities were then integrated using Weighted Nearest Neighbor analysis in Seurat. Graph-based Leiden clustering was performed on the weighted shared nearest-neighbour graph, and UMAP embeddings were generated for visualisation. The exact thresholds and parameters are provided on GitHub (https://github.com/Philpott-lab/SingleCellMultiOmics_Xenopus)

### Cell-type annotation

Cell clusters were annotated using RNA expression of established developmental marker genes previously described for *Xenopus Tropicalis atlas* (Briggs et al 2018). Annotations were assigned to major embryonic lineages and cell types where marker expression was sufficiently clear.

### Cell-type-specific differential expression analysis

Differential gene expression analysis was performed separately within each annotated cell type by comparing ASCL1-GR-treated cells with matched GFP control cells at each timepoint. For each comparison, cells from the relevant cell type and treatment/timepoint conditions were subsetted, and differential expression was performed using Seurat FindMarkers and Wilcoxon rank-sum test on SCT-normalised expression values. The sample condition was used as the identity class (ASCL1-GR-treated cells set as ident.1 and matched GFP control cells set as ident.2). Genes with positive log-fold change and adjusted p-value below the selected significance threshold were considered candidate ASCL1-upregulated genes. These upregulated genes were used for downstream Gene Ontology enrichment, target-gene classification and ChIP-linked accessibility analyses.

Differential expression testing was initially performed with logfc.threshold = 0.25 and min.pct = 0.1. For downstream classification of neuroectoderm ASCL1 response genes, genes were retained as induced targets if they showed positive differential expression with adjusted p-value < 0.05 and log2 fold-change > 1.

### Classification of ASCL1-induced and resistant target genes

Neuroectoderm ASCL1 response genes were defined as genes significantly upregulated in neuroectoderm after ASCL1 activation compared with control wild-type cells. For each non-neuroectodermal cell type, these neuroectoderm response genes were classified according to whether they were also significantly upregulated in that cell type or remained non-induced. Genes induced in both neuroectoderm and the comparison tissue were classified as shared or induced in that tissue, whereas genes induced in neuroectoderm but not in the comparison tissue were classified as resistant in that tissue.(Fig. 3).

### GO enrichment analysis

For Gene Ontology enrichment analysis, differentially upregulated Xenopus laevis genes were converted to human orthologs using a custom Xenopus–human–mouse ortholog conversion table, because functional annotation is more complete for human genes. Genes with valid human Entrez identifiers were retained for GO analysis. GO Biological Process enrichment was performed using clusterProfiler v4.14.6 and org.Hs.eg.db v3.20.0 with Benjamini–Hochberg multiple-testing correction. Enrichment was performed separately for each gene class or cell-type-specific upregulated gene set using enrichGO, with p-value cutoff 0.05 and q-value cutoff 0.2. Enriched terms were ranked by adjusted p-value.

### Assignment of ASCL1 ChIP-seq peaks to response genes

To link ASCL1 binding sites to candidate response genes, gene-associated regulatory regions were defined as the 50 kb region upstream of the transcription start site together with intronic regions of each gene. High-confidence ASCL1 ChIP-seq peaks (see section: ChIP data processing) overlapping these gene-associated regions were assigned to the corresponding gene. These ChIP-defined gene-linked regions were used for downstream snATAC-seq accessibility heatmaps, ATAC fragment quantification and motif analyses.

### snATAC-seq accessibility heatmaps at ChIP-defined ASCL1 sites

For region-level accessibility analysis, snATAC-seq fragments from the integrated Seurat/Signac object were quantified using ChIP-defined ASCL1 peaks linked to induced or resistant neuroectoderm response genes in other tissues/mesoderm. Region heatmaps for visualising ATAC fragment density were generated using Signac by centring ±10 kb window on each ChIP-defined ASCL1 defined candidate regulatory regions. Accessibility was compared between wild type control and ASCL1-treated nuclei within matched cell-type annotations and timepoints.

### HOMER motif enrichment analysis

HOMER motif enrichment analysis was performed on ChIP-defined ASCL1 candidate regulatory regions associated with neuroectoderm-induced genes at 1.25hr and 2.5hr after ASCL1 activation. Input regions were provided as BED files and analysed using findMotifsGenome.pl against the Xenopus laevis NCBI RefSeq genome assembly GCF_017654675.1. Motif enrichment was performed using the exact input region sizes (-size given), allowing up to two mismatches in motif optimisation (-mis 2) and reporting the top 25 motifs (-S 25). Known motif enrichment results from the HOMER output were used for downstream interpretation and Fig. generation.

### ChromVAR motif-associated accessibility analysis

For chromVAR analysis (Schep et al., 2017), snATAC-seq fragments were quantified within ChIP-defined ASCL1 peaks associated with neuroectoderm response genes using FeatureMatrix in Signac v1.14.0, and the resulting matrix was added to the Seurat v5.3.0/SeuratObject v5.1.0 object as a focused chromatin accessibility assay. Gene annotations for the *Xenopus laevis* NCBI RefSeq assembly GCF_017654675.1 were imported from the corresponding GTF file using rtracklayer v1.66.0. Annotation fields were reformatted to match the column names required by Signac, and protein-coding gene annotations were assigned to the ATAC assay before downstream chromatin accessibility analyses.

Motif matches within ChIP-defined ASCL1 regions were identified using vertebrate transcription factor position frequency matrices from JASPAR2022 v0.99.8 with TFBSTools v1.44.0. chromVAR deviation scores were then computed using Signac/chromVAR. These deviation scores estimate whether regions containing a given motif are more or less accessible than expected after correction for technical biases such as GC content.

To reduce sparsity in this focused ChIP-defined peak set, cells from the two ASCL1-activated timepoints, +ASCL1 1.25hr and +ASCL1 2.5hr, were combined before differential motif-associated accessibility analysis. Motifs with missing chromVAR deviation values in more than 20% of cells were excluded before downstream comparisons. Differential motif-associated accessibility was assessed using Seurat FindMarkers on the chromVAR assay, with cell type used as the identity class. Neuroectoderm was compared separately against endoderm, mesoderm, non-neuroectoderm, lateral plate mesoderm and epithelial skin. Testing was performed using logistic regression (test.use = “LR”) with min.pct = 0.1. Differential motif results were summarised using p-values, adjusted p-values, average log2 fold change and the percentage of cells with detectable motif deviation signal in each comparison group. Motifs with p-value < 0.05 were retained for exploratory downstream visualisation and summary tables, while adjusted p-values were also reported to assess multiple-testing-corrected significance.

### Figure illustrations

Icons and symbols to depict the different cell types and gene target groups were created using BioRender (BioRender.com). Drawings of *Xenopus* developmental stages were taken from Nieuwkoop and Faber (1994).

### Code availability and reproducibility

The main analysis R Markdown report is available on GitHub: https://github.com/Philpott-lab/SingleCellMultiOmics_Xenopus. It provides all input files, executable code, package requirements, analysis outputs, and automatically generated figures, enabling reproduction of the analyses presented in the manuscript.

