## Supplementary material for "Tissue-specific chromatin accessibility and co-factor availability together define ASCL1-dependent neural reprogrammability across germ layers during embryogenesis": SingleCellMultiOmics_Xenopus_analysis: SingleCellMultiOmics_Xenopus_analysis.html

Single-cell multi-omics analysis reproducible pipeline


Code 

- Show All Code
- Hide All Code

### Single-cell multi-omics analysis reproducible pipeline

##### Tissue-specific chromatin accessibility and cofactor availability together define Ascl1-dependent neural reprogrammability across germ layers during embryogenesis

###### David Lando^1,\*^, Samina Kausar^1,\*^, Toshiaki Shigeoka^1,2^, Frances Connor^1^, Jerome Jullien^3^ and Anna Philpott^1,4^ ^1^ Cambridge Stem Cell Institute, University of Cambridge, Cambridge CB2 0AW, UK ^2^ Division of Biological Science, Nara Institute of Science and Technology (NAIST), Ikoma-shi, Nara, Japan ^3^ Center for Research in Transplantation and Translational Immunology, University of Nantes, Nantes, 44000, France ^4^ Department of Oncology, University of Cambridge, Cambridge CB2 0AH, UK \* These authors contributed equally to this work ^4^ To whom correspondence should be addressed

#### 2026-07-07

- 1 Overview
- 2 Package loading
- 3 Metadata, parameters and helper
  functions
- 4 Global settings and file
  paths
- 5 Load raw snRNA-seq data and build
  merged Seurat object
  - 5.1 RNA quality control
- 6 Single Cell Multi-Omics joint
  analysis
  - 6.1 Match RNA cells to ATAC fragment
    barcodes
  - 6.2 Create filtered fragment
    object
  - 6.3 Build gene-structure-window ATAC
    feature set
  - 6.4 Create chromatin assay and ATAC QC
    metrics
  - 6.5 ATAC filtering
  - 6.6 RNA and ATAC dimensional reduction
    for multimodal integration
  - 6.7 WNN integration and
    clustering
  - 6.8 Manual cluster annotation
- 7 Differential expression analysis: a
  template for defining neuroectoderm-induced genes
  - 7.1 DE analysis within the same cell
    type across conditions +Ascl1\_T1\_25 vs. WT\_T1\_25
  - 7.2 GO enrichment analysis
- 8 Ascl1 Binding sites (MACS peaks)
  - 8.1 Ascl1 Binding sites regionMatrix
    to visualise chromatin accessibility: inducible and non-inducible genes
    scATAC heatmaps
  - 8.2 Ascl1 Binding sites boxplots to
    compare chromatin accessibility
- 9 Motif and chromatin accessibility
  analysis to identifies potential signatures associated with Ascl1 gene
  regulation
  - 9.1 HOMER Analysis
  - 9.2 chromVAR
    - 9.2.1 scATAC using Ascl1 Chipseq.
      peaks of selected list of response genes
    - 9.2.2 Compute motifs deviations scores
      in ChromVAR assay
    - 9.2.3 Find differentially accessible
      motifs between cell types (+Ascl1\_T1\_25 and +Ascl1\_T2\_5)
    - 9.2.4 Top Motifs from HOMER and
      chromVAR
- 10 Sox motif family expression data
  visualization
- 11 Session information

### 1 Overview

This report contains the R Markdown workflow used to analyse
single-cell multiomics and related genomic datasets for the study of
Ascl1-dependent neural reprogramming in the developing *Xenopus
laevis* embryo. The report includes executable code, package
requirements, analysis outputs, and automatically generated figures
associated with the manuscript.

The goal of this report is to provide the key analysis steps required
to reproduce the main results. Code for these important steps is
included directly within the report.

### 2 Package loading

```
library(Seurat)
library(SeuratObject)
library(Signac)
library(ggplot2)
library(dplyr)
library(stringr)
library(patchwork)
library(rtracklayer)
library(GenomicRanges)
library(Matrix)
library(future)
library(DT)
library(profileplyr)
library(DiffBind)
library(ChIPseeker)
library(GenomicFeatures)
library(rvest)
library(tidyr)
library(purrr)
library(rvest)
library(pheatmap)
library(universalmotif)
library(ggseqlogo)
library(cowplot)
library(Rsamtools)
library(TFBSTools)
library(JASPAR2022)
library(ggtext)

library(tibble)
library(clusterProfiler)
library(org.Hs.eg.db)
library(org.Mm.eg.db)
library(enrichplot)
library(tidytext)
```

### 3 Metadata, parameters and helper functions

```
#-------------------------------
# Samples
#-------------------------------
sample_names <- c(
  "DNAA001","DNAA002","DNAA003","DNAA004","DNAA005","DNAA006",
  "DNAA007","DNAA008","DNAA009","DNAA010","DNAA011","DNAA012",
  "DNAA014","DNAA015","DNAA016","DNAA017","DNAA018","DNAA019",
  "DNAA020","DNAA021","DNAA022","DNAA023"
)

sample_ids <- c(
  "01", "02", "03", "04", "05", "06",
  "07", "08", "09", "10", "11", "12",
  "14", "15", "16", "17", "18", "19",
  "20", "21", "22", "23"
)

#-------------------------------
# Metadata mapping
#-------------------------------
replicate_map <- c(
  "01" = "GFPT0r1", "02" = "GFPT0r2",
  "03" = "AsclT0r1", "04" = "AsclT0r2",
  "05" = "GFPT1_25r1", "06" = "GFPT1_25r2",
  "07" = "AsclT1_25r1", "08" = "AsclT1_25r2",
  "09" = "GFPT2_5r1", "10" = "GFPT2_5r2",
  "11" = "AsclT2_5r1", "12" = "AsclT2_5r2"
)

condition_map <- c(
  "01" = "WT_T0",      "02" = "WT_T0",
  "03" = "+Ascl1_T0", "04" = "+Ascl1_T0",
  "05" = "WT_T1_25",  "06" = "WT_T1_25",
  "07" = "+Ascl1_T1_25", "08" = "+Ascl1_T1_25",
  "09" = "WT_T2_5",   "10" = "WT_T2_5",
  "11" = "+Ascl1_T2_5", "12" = "+Ascl1_T2_5"
)

#-------------------------------
# Filtering / analysis parameters
#-------------------------------
rna_min_cells <- 3
rna_min_features <- 200
rna_min_features_QC <- 500
rna_max_mito_QC <- 5

atac_fragment_cutoff <- 1000
atac_min_features <- 100
atac_min_cells_feature <- 5

n_variable_features <- 2000
n_pcs_rna <- 50
n_lsi <- 100
umap_dims_rna <- 1:12
wnn_rna_dims <- 1:50
wnn_atac_dims <- 2:100
cluster_resolution_rna <- 0.2
cluster_resolution_wnn <- 0.1

################################################################# functions
make_seurat_object <- function(counts_mat, project_name) {
  obj <- CreateSeuratObject(
    counts = counts_mat,
    project = project_name,
    min.cells = rna_min_cells,
    min.features = rna_min_features
  )

  # Cell barcode format expected to contain sample ID after ':'
  obj$sampleID <- str_extract(colnames(obj), "(?<=:)[^:]+$")
  obj$sample_name <- unname(replicate_map[obj$sampleID])
  obj$sample_condition <- unname(condition_map[obj$sampleID])
  obj$lib <- as.character(obj$orig.ident)

  obj
}

add_batch_label <- function(seurat_list) {
  for (i in seq_along(seurat_list)) {
    seurat_list[[i]]$Dataset <- ifelse(i <= 10, "Batch1", "Batch2")
  }
  seurat_list
}


# Function for data parsing: Homer analysis
parse_homer_html <- function(html_file, cell_type, motif_type, top_n = NULL) {
  if (!file.exists(html_file)) return(NULL)
  
  page <- read_html(html_file)
  tab <- html_table(html_nodes(page, "table")[[1]], fill = TRUE)
  df <- as.data.frame(tab, stringsAsFactors = FALSE)
  
  if (motif_type == "known") {
    colnames(df) <- c("Rank", "Motif", "Name", "P_value", "log_P_pvalue", "q_value",
                      "Target_with_motif", "Target_percent", 
                      "Background_with_motif", "Background_percent", "Motif_File", "SVG")
    
    df <- df %>%
      filter(!Name %in% c("Name")) %>%  # Remove header row
      mutate(
        Rank= as.numeric(Rank),
        P_value = as.numeric(P_value),
        P_log10 = -log10(P_value),
        Target_percent = as.numeric(str_remove(Target_percent, "%")),
        Background_percent = as.numeric(str_remove(Background_percent, "%")),
        Cell_Type = cell_type,
        Motif_Type = motif_type
      ) %>%
       dplyr::select(Rank, Cell_Type, Motif_Type, Name, P_value, P_log10, Target_percent, Background_percent)
    
  } else if (motif_type == "denovo") {
    colnames(df) <- c("Rank", "Motif", "P_value", "log_P_pvalue", "Target_percent", 
                      "Background_percent", "STD", "Best_Match", "Motif_File")
    
    df <- df %>%
      filter(!Best_Match %in% c("Best Match/Details")) %>%
      mutate(
        P_value = as.numeric(P_value),
        P_log10 = -log10(P_value),
        Target_percent = as.numeric(str_remove(Target_percent, "%")),
        Background_percent = as.numeric(str_remove(Background_percent, "%")),
        Cell_Type = cell_type,
        Motif_Type = motif_type
      ) %>%
      select(Cell_Type, Motif_Type, Best_Match, P_value, P_log10, Target_percent, Background_percent) %>%
      rename(Name = Best_Match)
  } else {
    return(NULL)
  }
  
  # Optional: only keep top N if requested
  if (!is.null(top_n)) {
    df <- df %>% slice_head(n = top_n)
  }
  
  return(df)
}
```

### 4 Global settings and file paths

Replace these paths with your local project paths before running.

### 5 Load raw snRNA-seq data and build merged Seurat object

The single-cell multiomics data were initially generated from
split-pool barcoding libraries. Preprocessing of the split-pool data was
performed using the Paired-Tag processing workflow (https://github.com/cxzhu/Paired-Tag), which produces
multiple intermediate files for RNA and ATAC modalities across
sublibraries.

For the analyses in this report, we started from the original
preprocessed RNA count files generated for each sublibrary. Initial
barcode and count-based filtering steps were applied, after which all
sublibraries were merged to generate a single combined snRNA-seq gene
count matrix for downstream analysis in Seurat.

For GEO submission, we provided the filtered and merged files
generated after these initial processing steps. For the chromatin
accessibility modality, the filtered scATAC fragment file was submitted
as: **sc\_multiome\_ATAC\_filtered.tbi and .tsv**

For the transcriptomic modality, a **merged scRNA/snRNA-seq
gene count matrix** is provided. Therefore, although the GEO
submission contains filtered and merged files, this section documents
the analysis workflow starting from the original preprocessed
sublibrary-level files before merging and downstream Seurat object
construction.

```
loaded_data_list <- list()

for (nm in sample_names) {
  sample_dir <- file.path(rna_10x_dir, nm)
  if (!file.exists(sample_dir)) next
  loaded_data_list[[nm]] <- Read10X(data.dir = sample_dir)
}

stopifnot(length(loaded_data_list) > 0)

rna_list <- lapply(names(loaded_data_list), function(nm) {
  make_seurat_object(loaded_data_list[[nm]], project_name = nm)
})
names(rna_list) <- names(loaded_data_list)

rna_list <- add_batch_label(rna_list)

rna_obj <- merge(
  x = rna_list[[1]],
  y = rna_list[-1],
  add.cell.ids = sample_ids[seq_along(rna_list)]
)

# Seurat v5 layer joining
rna_obj[["RNA"]] <- JoinLayers(rna_obj[["RNA"]])

# Harmonize barcode format with fragment file format
new_barcodes <- gsub("_", ":", colnames(rna_obj))
colnames(rna_obj) <- new_barcodes
rownames(rna_obj@meta.data) <- new_barcodes

rna_obj$numID <- seq_len(ncol(rna_obj))

rna_obj
```

```
## An object of class Seurat 
## 39023 features across 75073 samples within 1 assay 
## Active assay: RNA (39023 features, 0 variable features)
##  1 layer present: counts
```

#### 5.1 RNA quality control

```
# Mitochondrial genes in Xenopus may need dataset-specific curation.
# Adjust the pattern if your annotation uses a different naming convention.
#mito_features <- grep("^MT-|^mt-", rownames(rna_obj), value = TRUE)

#######################################################################
########################## surat 5 add pattern for Xenopus v10 MT genes as they do not have any pattern in names
###....;;;;;;;;;;;;;;;;;;;;;;;;;;;;;;;;;;;;;;;;;;

# Step 1: Extract the count matrix from the Seurat object
# Step 1: Define mitochondrial gene names
mt_gene_names <- c("ND1", "ND2", "COX1", "COX2", "ATP8", "ATP6", 
                   "COX3", "ND3", "ND4L", "ND4", "ND5", "ND6", "CYTB")

# Step 2: Get the current feature names
original_gene_names <- rownames(rna_obj[["RNA"]])

# Step 3: Add "mt-" prefix for mitochondrial genes
new_gene_names <- ifelse(original_gene_names %in% mt_gene_names, 
                         paste0("mt-", original_gene_names), 
                         original_gene_names)

# Step 4: Directly update feature names in the assay
rownames(rna_obj[["RNA"]]) <- new_gene_names

# Step 5: Verify the update
head(rownames(rna_obj[["RNA"]]))
```

```
## [1] "1a11.L"   "42sp43.L" "42sp50.L" "mt-ATP6"  "mt-ATP8"  "mt-COX1"
```

```
# Step 6: Calculate mitochondrial percentages
mito_genes <- grep("^mt-", new_gene_names, value = TRUE, ignore.case = TRUE)
rna_obj$percent.mito <- PercentageFeatureSet(rna_obj, features = mito_genes)

# View percentage statistics
summary(rna_obj$percent.mito)
```

```
##    Min. 1st Qu.  Median    Mean 3rd Qu.    Max. 
##  0.0000  0.0000  0.2530  0.4055  0.5089 10.3870
```

```
#;;;;;;;;;;;;;;;;;;;;;;;;;;;;;;;;;;;;;;;;;;;;;;;

if (length(mito_genes) > 0) {
  rna_obj[["percent.mito"]] <- PercentageFeatureSet(rna_obj, features = mito_genes)
} else {
  rna_obj[["percent.mito"]] <- 0
}

VlnPlot(
  rna_obj,
  features = c("nCount_RNA", "nFeature_RNA", "percent.mito"),
  group.by = "sample_condition",
  pt.size = 0.05,
  log = TRUE,
  ncol = 3
)
```

```
# QC filtering using metadata columns directly (robust in Seurat v5)
keep_cells <- with(
  rna_obj@meta.data,
  percent.mito < rna_max_mito_QC &
  nFeature_RNA > rna_min_features_QC
)

rna_obj <- rna_obj[, keep_cells]

rna_obj
```

```
## An object of class Seurat 
## 39023 features across 38208 samples within 1 assay 
## Active assay: RNA (39023 features, 0 variable features)
##  1 layer present: counts
```

```
# verify seurat object that it has the right number of cells after all filtering steps
datatable(data.frame((table(rna_obj$sample_name))), 
          colnames = c("Sample","Nb of cells"),
          rownames = FALSE)
```

```
# verify seurat object that it has the right number of cells after all filtering steps
datatable(data.frame((table(rna_obj$sample_condition))), 
          colnames = c("Conditions","Nb of cells"),
          rownames = FALSE)
```

```
# verify seurat object that it has the right number of cells after all filtering steps
datatable(data.frame((table(rna_obj$lib))), 
          colnames = c("Lib","Nb of cells"),
          rownames = FALSE)
```

```
#Save RNA-only checkpoint
#saveRDS(rna_obj, file.path(output_dir, "scRNA_seq_object_afterQCfilters.rds"))
```

### 6 Single Cell Multi-Omics joint analysis

#### 6.1 Match RNA cells to ATAC fragment barcodes

```
# Count fragments from merged fragment file
fragment_counts <- CountFragments(merged_fragment_file)
selected_barcodes <- fragment_counts$CB[fragment_counts$frequency_count > atac_fragment_cutoff]

common_barcodes <- intersect(colnames(rna_obj), selected_barcodes)
length(common_barcodes)
```

```
## [1] 24282
```

```
multiome_obj <- subset(rna_obj, cells = common_barcodes)
multiome_obj
```

```
## An object of class Seurat 
## 39023 features across 24282 samples within 1 assay 
## Active assay: RNA (39023 features, 0 variable features)
##  1 layer present: counts
```

#### 6.2 Create filtered fragment object

For strict reproducibility, we have generated and archived the
filtered fragment file once which is available on GEO.

```
#filtered_fragment_file <- file.path(output_dir, "FragmentFile_filtered_barcodes_cutoff1000_bothBatches.tsv.gz")

#FilterCells(
#  fragments = merged_fragment_file,
#  cells = common_barcodes,
#  outfile = filtered_fragment_file,
#  buffer_length = 256L,
#  verbose = TRUE
#)
```

```
#filtered_fragment_file <- file.path(output_dir, "FragmentFile_filtered_barcodes_cutoff1000_bothBatches.tsv.gz")

frags <- CreateFragmentObject(
  path = filtered_fragment_file,
  cells = common_barcodes
)
```

#### 6.3 Build gene-structure-window ATAC feature set

For the ATAC modality, we evaluated alternative feature definitions
because peak-based feature construction from shallow split-pool
scATAC-seq data in Xenopus laevis produced sparse and less stable
matrices. We compared standard peak-based features, fixed-width genomic
bins and gene-structure-window features. Gene-structure-window features
were defined using annotated gene bodies together with upstream
regulatory windows, generating an interpretable feature space linked to
gene structure. Feature definitions were benchmarked by comparing
ATAC-derived embeddings and clusters against RNA-derived cell-type
annotations from the paired multiome data. Gene-structure-window
features showed improved agreement with RNA-defined cell identities
compared with peak-based features and were therefore selected for the
final ATAC feature matrix for downstream dimensionality reduction and
multiome integration.

```
gtf <- import(gtf_file)
# For Xenopus genome: rename column with standered required names
colnames(mcols(gtf))[colnames(mcols(gtf)) == "gene"] <- "gene_name"
colnames(mcols(gtf))[colnames(mcols(gtf)) == "transcript_id"] <- "tx_id"

protein_coding_genes <- gtf[!is.na(gtf$gene_biotype) & gtf$gene_biotype == "protein_coding"]

# Keep only protein coding genes
pc_genes <- gtf[gtf$type == "gene" & gtf$gene_biotype == "protein_coding"]
# Keep only exons belonging to protein coding genes
pc_exons <- gtf[gtf$type == "exon" & gtf$gene_id %in% pc_genes$gene_id]

# Split gene bodies and exons by gene_id
genes_by_id <- split(pc_genes, pc_genes$gene_id)
exons_by_id <- GenomicRanges::reduce(split(pc_exons, pc_exons$gene_id)) 

exons <- unlist(exons_by_id)
exons$type <- "exon"
exons$gene_id <- rep(names(exons_by_id), lengths(exons_by_id))
exons <- exons[width(exons) >= 50 & width(exons) <= 5e4]

introns <- readRDS(introns_rds) #      <-------------------    its just getting introns coordinates (file: protein_coding_introns.rds) generated from gtf
introns <- introns[width(introns) >= 50]

upstream <- promoters(pc_genes, upstream = 5000, downstream = 0)
start(upstream)[start(upstream) < 1] <- 1
upstream$type <- "Upstream5kb"

# Final ATAC feature set used in the original pipeline:
# upstream promoter windows + exons + introns
upstream_exons_introns <- c(upstream, exons, introns)
```

#### 6.4 Create chromatin assay and ATAC QC metrics

```
atac_counts <- FeatureMatrix(
  fragments = frags,
  features = upstream_exons_introns,
  cells = common_barcodes
)

# Ensure order matches Seurat object
atac_counts <- atac_counts[, colnames(multiome_obj)]

multiome_obj[["ATAC"]] <- CreateChromatinAssay(
  counts = atac_counts,
  sep = c(":", "-"),
  fragments = filtered_fragment_file,
  min.cells = atac_min_cells_feature,
  min.features = 0
)

# Remove cells lacking valid ATAC assay values after assay creation
DefaultAssay(multiome_obj) <- "RNA"
multiome_obj <- subset(
  multiome_obj,
  cells = rownames(multiome_obj@meta.data)[!is.na(multiome_obj$nCount_ATAC)]
)

# Add fragment count metadata
fragment_counts_subset <- fragment_counts[fragment_counts$CB %in% colnames(multiome_obj), ]
multiome_obj$frequency_count <- fragment_counts_subset$frequency_count[
  match(colnames(multiome_obj), fragment_counts_subset$CB)
]

# Signac QC metrics
DefaultAssay(multiome_obj)<- "ATAC"
Annotation(multiome_obj) <- protein_coding_genes
multiome_obj <- FRiP(multiome_obj, assay = "ATAC", total.fragments = "frequency_count")
multiome_obj <- TSSEnrichment(multiome_obj, region_extension = 2000)
multiome_obj
```

```
## An object of class Seurat 
## 482915 features across 24282 samples within 2 assays 
## Active assay: ATAC (443892 features, 0 variable features)
##  2 layers present: counts, data
##  1 other assay present: RNA
```

#### 6.5 ATAC filtering

```
VlnPlot(
  multiome_obj,
  features = c("nCount_ATAC", "nFeature_ATAC", "frequency_count", "FRiP", "TSS.enrichment"),
  group.by = "sample_condition",
  pt.size = 0.05,
  ncol = 5
)
```

```
# -----------------------------
# ATAC QC filtering
# -----------------------------
keep_cells <- with(
  multiome_obj@meta.data,
  nCount_ATAC > 0 &
  nFeature_ATAC > atac_min_features
)

multiome_obj <- multiome_obj[, keep_cells]


DensityScatter(
  multiome_obj,
  x = "nFeature_ATAC",
  y = "TSS.enrichment",
  log_x = TRUE,
  quantiles = TRUE
)
```

#### 6.6 RNA and ATAC dimensional reduction for multimodal integration

```
# RNA modality
DefaultAssay(multiome_obj) <- "RNA"
multiome_obj <- SCTransform(
  multiome_obj,
  assay = "RNA",
  variable.features.n = n_variable_features,
  verbose = FALSE
)

DefaultAssay(multiome_obj) <- "SCT"
multiome_obj <- RunPCA(multiome_obj, assay = "SCT", npcs = n_pcs_rna, verbose = FALSE)

# ATAC modality
DefaultAssay(multiome_obj) <- "ATAC"
multiome_obj <- RunTFIDF(multiome_obj, assay = "ATAC")
multiome_obj <- FindTopFeatures(multiome_obj, min.cutoff = 15, assay = "ATAC")
multiome_obj <- RunSVD(multiome_obj, assay = "ATAC", n = n_lsi)

ElbowPlot(multiome_obj, reduction = "pca", ndims = 50)
```

```
DepthCor(multiome_obj,  assay = "ATAC")
```

#### 6.7 WNN integration and clustering

```
multiome_obj <- FindMultiModalNeighbors(
  multiome_obj,
  reduction.list = list("pca", "lsi"),
  dims.list = list(wnn_rna_dims, wnn_atac_dims)
)

multiome_obj <- RunUMAP(
  multiome_obj,
  nn.name = "weighted.nn",
  reduction.name = "wnn.umap",
  reduction.key = "wnnUMAP_"
)

multiome_obj <- FindClusters(
  multiome_obj,
  graph.name = "wsnn",
  algorithm = 4,
  resolution = cluster_resolution_wnn,
  verbose = FALSE
  , random.seed = 0
)

# Modality-specific UMAPs for comparison
DefaultAssay(multiome_obj) <- "SCT"
multiome_obj <- RunUMAP(
  multiome_obj,
  dims = wnn_rna_dims,
  reduction.name = "umap.rna",
  reduction.key = "rnaUMAP_"
)

DefaultAssay(multiome_obj) <- "ATAC"
multiome_obj <- RunUMAP(
  multiome_obj,
  reduction = "lsi",
  dims = wnn_atac_dims,
  reduction.name = "umap.atac",
  reduction.key = "atacUMAP_"
)

Idents(multiome_obj) <- multiome_obj$wsnn_res.0.1

p_rna <- DimPlot(multiome_obj, reduction = "umap.rna", label = TRUE) + ggtitle("RNA")
p_atac <- DimPlot(multiome_obj, reduction = "umap.atac", label = TRUE) + ggtitle("ATAC")
p_wnn <- DimPlot(multiome_obj, reduction = "wnn.umap", label = TRUE) + ggtitle("WNN")

p_rna + p_atac + p_wnn
```

#### 6.8 Manual cluster annotation

We used marker genes and domain knowledge to annotate clusters. The
labels below reflect the mapping in the original analysis and should be
verified against the final chosen clustering.

```
Idents(multiome_obj) <- multiome_obj$seurat_clusters

new.cluster.ids <- c(
  
  "Neuroectoderm",
  "Mesoderm",
  "Endoderm",
  "Lateral-plate-mesoderm",
  "Epithelial-skin",
  "Non-neuroectoderm",
  "Fibroblast",
  "Ciliated-epidermal-skin",
  "9",
  "10"
)

if (length(new.cluster.ids) == length(levels(multiome_obj))) {
  names(new.cluster.ids) <- levels(multiome_obj)
  multiome_obj <- RenameIdents(multiome_obj, new.cluster.ids)
  multiome_obj$cell_type <- Idents(multiome_obj)
}

marker_features <- c( "meox2.L","myh4.L", "des.1.L", "sox3.S", "mapt.S", "dcc.S","darmin.L","hand1.L", "epcam.L", "tp63.S", "krt12.4.S", "krt70.L",     "emilin3.L", "col8a1.L", "dnah9.S", "foxj1.S")

my_cols <- c(
"Mesoderm" = "#E7298A",
"Neuroectoderm" = "royalblue1",
"Endoderm" = "mediumpurple",
"Lateral-plate-mesoderm" = "pink2",
"Epithelial-skin" = "skyblue4",
"Non-neuroectoderm" = "#E6AB02",
"Fibroblast" = "#D95F02",
"Ciliated-epidermal-skin" = "cyan2",
"9" = "honeydew3",
"10" = "snow4")

my_cols2 <- c(
"WT_T0" = "lightblue",
"WT_T1_25" = "royalblue1",
"WT_T2_5" = "darkblue",
"+Ascl1_T0" = "pink",
"+Ascl1_T1_25" = "salmon2",
"+Ascl1_T2_5" = "red3"
)
# ........................................................................ Figure 1 Study of Ascl1 directed reprograming in Xenopus development. 

DefaultAssay(multiome_obj) <- "SCT"
DimPlot(multiome_obj, reduction = "wnn.umap", group.by = "cell_type", label = TRUE) +
          scale_color_manual(values = my_cols)
```

```
DimPlot(multiome_obj, reduction = "wnn.umap",  group.by = "sample_condition"  )  +
          scale_color_manual(values = my_cols2)
```

```
DotPlot(multiome_obj, features = marker_features, assay = "SCT") + RotatedAxis() +
  scale_color_gradientn(colors = c(
    "#FDE0DD",
    "#FA9FB5",
    "#C51B8A",
    "#7A0177"
  ))
```

```
#Save final integrated object
#saveRDS(multiome_obj, file.path(output_dir, "Xenopus_multiome_integrated_submission.rds"))
```

### 7 Differential expression analysis: a template for defining neuroectoderm-induced genes

#### 7.1 DE analysis within the same cell type across conditions +Ascl1\_T1\_25 vs. WT\_T1\_25

The original files contain extensive downstream DE. A minimal
reproducible template is shown below.

Genes with positive log-fold change and adjusted p-value below the
selected significance threshold were considered candidate
Ascl1-upregulated genes. These upregulated genes were used for
downstream GO enrichment, target-gene classification and ChIP-linked
accessibility analyses.

Differential expression testing was initially performed with
logfc.threshold = 0.25 and min.pct = 0.1. For downstream classification
of neuroectoderm Ascl1 response genes, genes were retained as induced
targets if they showed positive differential expression with adjusted
p-value < 0.05 and log2 fold-change > 1.

```
DefaultAssay(multiome_obj) <- "SCT"
library(future)
options(future.globals.maxSize = 8 * 1024^3)  # 8 GiB


# Example: compare +Ascl1_T1_25 vs WT_T1_25 within one cell type
sub_obj <- subset(
  multiome_obj,
  subset = cell_type == "Neuroectoderm" &
           sample_condition %in% c("WT_T1_25", "+Ascl1_T1_25")
)

Idents(sub_obj) <- sub_obj$sample_condition

markers_t1 <- FindMarkers(
  sub_obj,
  ident.1 = "+Ascl1_T1_25",
  ident.2 = "WT_T1_25",
  assay = "SCT",
  slot = "data",  # recomended for sct the corrected umi
  test.use = "wilcox",
  logfc.threshold = 0.25,
  min.pct = 0.1,
  verbose = FALSE)


head(markers_t1)
```

```
##                   p_val avg_log2FC pct.1 pct.2     p_val_adj
## myt1.S    1.281749e-158   2.646409 0.786 0.264 4.794640e-154
## myt1.L    8.436035e-119   2.173761 0.720 0.227 3.155668e-114
## dll1.L    1.455379e-113   2.589256 0.627 0.154 5.444135e-109
## zc3h12c.L  1.052077e-88   2.791137 0.509 0.107  3.935503e-84
## cbfa2t2.L  2.411927e-82   2.762993 0.477 0.094  9.022294e-78
## dll1.S     1.980587e-75   2.008422 0.546 0.164  7.408783e-71
```

```
#write.csv(markers_t1, file.path(output_dir, "DE_Neuroectoderm_Ascl1T1_25_vs_WT.csv"))
```

#### 7.2 GO enrichment analysis

For simplicity, only the GO analysis of the Neuroectoderm is shown in
this report.

For Gene Ontology enrichment analysis, differentially upregulated
Xenopus laevis genes were converted to human orthologs using a custom
Xenopus–human–mouse ortholog conversion table (provided on Github),
because functional annotation is more complete for human genes. Genes
with valid human Entrez identifiers were retained for GO analysis.

```
##          ID gene_short_name      gene_short_name_pre_release simple_name
## 1     gene1       slc26a4.L                        slc26a4.L     slc26a4
## 2    gene10    LOC108710558 LOC108710558 [provisional:wdr91]       wdr91
## 3   gene100         dhx16.S                          dhx16.S       dhx16
## 4  gene1000          ufm1.L                           ufm1.L        ufm1
## 5 gene10000     LOC495506.S                       cyp2c8.1.L    cyp2c8.1
## 6 gene10001        prdm12.L                         prdm12.L      prdm12
##               Xenabse human_entrez human_symbol human_symbol_refined
## 1                <NA>           NA         <NA>              SLC26A4
## 2  XB-GENEPAGE-969791        29062        WDR91                WDR91
## 3  XB-GENEPAGE-480524         8449        DHX16                DHX16
## 4  XB-GENEPAGE-973174        51569         UFM1                 UFM1
## 5 XB-GENEPAGE-5831570         1558       CYP2C8               CYP2C8
## 6  XB-GENEPAGE-953728        59335       PRDM12               PRDM12
##     human_Ensembl human_ENTREZ_refined Neuronal_genes      Mouse_Ensembl
## 1 ENSG00000091137                 5172              0 ENSMUSG00000020651
## 2 ENSG00000105875                29062              0 ENSMUSG00000058486
## 3 ENSG00000227222                 8449              0                  0
## 4 ENSG00000120686                51569              0 ENSMUSG00000027746
## 5 ENSG00000138115                 1558              0               <NA>
## 6 ENSG00000130711                59335              1 ENSMUSG00000079466
##   Mouse_gene Mouse_gene_refined
## 1    Slc26a4            Slc26a4
## 2      Wdr91              Wdr91
## 3          0              Dhx16
## 4       Ufm1               Ufm1
## 5       <NA>             Cyp2c8
## 6     Prdm12             Prdm12
```

```
## [1] "Top DEGs for each cluster: Input for mapping orthologues"
```

```
## $Neuroectoderm
##   [1] "myt1.S"       "myt1.L"       "dll1.L"       "zc3h12c.L"    "cbfa2t2.L"   
##   [6] "dll1.S"       "spsb4.L"      "cbfa2t2.S"    "LOC108703481" "pak3.S"      
##  [11] "celf3.L"      "zc3h12c.S"    "rbm24.L"      "tspan33.L"    "celf2.L"     
##  [16] "nfasc.S"      "LOC108712923" "LOC108698041" "rbfox2.S"     "ebf2.S"      
##  [21] "amotl2.L"     "LOC108699846" "magi1.S"      "znf238.2.L"   "rnf165.L"    
##  [26] "smpd3.S"      "LOC121397066" "evi5.L"       "slc30a8.L"    "ulk1.S"      
##  [31] "dlc.L"        "ulk1.L"       "cdc25b.L"     "LOC108717058" "lmx1b.1.L"   
##  [36] "LOC108706545" "spsb4.S"      "ebf2.L"       "map3k13.L"    "notch1.S"    
##  [41] "LOC108711245" "LOC108709943" "stard13.S"    "neurl1b.L"    "elavl3.L"    
##  [46] "pag1.L"       "dio2.L"       "itga6.L"      "rap1gap2.L"   "gadd45g.S"   
##  [51] "slc16a3.L"    "ncs1.S"       "ca8.L"        "gas6.L"       "cd82.S"      
##  [56] "tead4.L"      "rbfox2.L"     "tox3.L"       "srsf5.S"      "wnt11b.L"    
##  [61] "pak3.L"       "hes6.2.L"     "LOC108698411" "LOC108707564" "gse1.L"      
##  [66] "slco3a1.S"    "sertad2.L"    "LOC108704496" "LOC108714671" "LOC108716920"
##  [71] "b4galnt1.S"   "tead1.S"      "lmx1b.1.S"    "mob3b.S"      "rfx3.L"      
##  [76] "arhgap10.L"   "enox1.L"      "LOC108716929" "bin1.S"       "LOC108715881"
##  [81] "gse1.S"       "plk3.S"       "gadd45g.L"    "agmo.L"       "fam102a.L"   
##  [86] "LOC108701330" "pkp4.S"       "nol4.S"       "plxna2.L"     "znf238.2.S"  
##  [91] "mtcl1.L"      "LOC108698803" "ankrd35.L"    "onecut1.L"    "mxra7.L"     
##  [96] "fgfr4.L"      "myo6.L"       "traf4.S"      "mob2.2.L"     "kcnk9.L"     
## [101] "id3.S"        "hes6.1.S"     "rgma.L"       "LOC108710286" "galnt16.L"   
## [106] "thsd7a.S"     "mxi1.L"       "rnf165.S"     "runx1t1.L"    "wif1.L"      
## [111] "LOC108713414" "foxp2.L"      "n4bp2.L"      "igfbp5.S"     "smpd3.L"     
## [116] "LOC121394543" "abca3.L"      "LOC121401529" "antxr2.S"     "ptk2.L"      
## [121] "foxp2.S"      "amotl2.S"     "nol4.L"       "LOC121400519" "btc.L"       
## [126] "plk3.L"       "ddr1.L"       "lmo2.S"       "LOC108715931" "LOC108718775"
## [131] "LOC108702752" "LOC121393757" "LOC108719216" "plxna2.S"     "tacc1.L"     
## [136] "LOC121402966" "LOC121398761" "egfl6.L"      "kiaa1217.S"   "arid1b.L"    
## [141] "LOC121393708" "igdcc3.L"     "LOC108707093" "kif5c.S"      "LOC108698571"
## [146] "cdknx.L"      "trim2.S"      "LOC121403010" "abca3.S"      "nfe2l3.S"    
## [151] "LOC108715172" "dbn1.S"       "lrrn1.S"      "prox1.S"      "celf3.S"     
## [156] "LOC108711262" "fam102a.S"    "thsd7a.L"     "LOC121399491" "btc.S"       
## [161] "bcr.L"        "pdk4.S"       "akt1.S"       "rbm38.L"      "LOC121395456"
## [166] "slc7a6.L"     "prickle2.L"   "map3k13.S"    "LOC121393758" "myo5b.L"     
## [171] "notch1.L"     "frmpd1.L"     "LOC108697535" "LOC108699933" "LOC121396893"
## [176] "LOC121401562" "onecut2.S"    "LOC108705236" "myo10.2.S"    "cldn6.2.L"   
## [181] "LOC108698659" "LOC121401551" "dbn1.L"       "LOC108699934" "ebf3.S"      
## [186] "hes5.5.L"     "LOC108715723" "fryl.S"       "c2cd2.S"      "rbm38.S"     
## [191] "wtip.S"       "pcsk5.S"      "LOC108699342" "LOC121400501" "LOC108717893"
## [196] "id3.L"        "onecut2.L"    "cd82.L"       "dcc.S"        "LOC121394660"
## [201] "mb21d2.L"     "tead4.S"      "vash2.L"      "LOC121396258" "kiaa1217.L"  
## [206] "ptbp3.L"      "cntrl.L"      "pcbp2.S"      "mb21d2.S"     "pnrc2.S"     
## [211] "kcnn3.L"      "LOC108709567" "gprc5cl1.L"   "tfdp2.L"      "rasgef1b.L"  
## [216] "plin3.S"      "enah.L"       "pcdh7.L"      "LOC108706611" "LOC108710980"
## [221] "rnf43.S"      "LOC121400500" "rbm24.S"      "lrrn1.L"      "LOC121393376"
## [226] "LOC121397070" "cldn6.2.S"    "pkig.L"       "lrp4.S"       "tspan9.L"    
## [231] "LOC121402983" "post.L"       "LOC108699920" "rnf43.L"      "ywhaz.S"     
## [236] "nfe2l3.L"     "LOC108695382" "LOC108715740" "skp1.L"       "c1orf21.S"   
## [241] "LOC108719816" "grip1.S"      "LOC108716472" "LOC108698619" "ror2.L"      
## [246] "LOC121402182" "meox2.L"      "LOC108698445" "ada2.L"       "hes5.1.S"    
## [251] "LOC108715827" "pcdh18.L"     "LOC108710721" "pknox2.L"     "tshz1.S"     
## [256] "magi1.L"      "bcr.S"        "onecut1.2.S"  "lrrc8e.L"     "xicl"        
## [261] "spry4.L"      "LOC108701655" "LOC108712669" "nin.L"        "XB22041735.L"
## [266] "plekhg4.L"
```

```
##  [1] "cbfa2t2.S"    "LOC108698041" "LOC121397066" "LOC108706545" "rbfox2.L"    
##  [6] "LOC121394543" "LOC121401529" "LOC121400519" "LOC108702752" "LOC121393757"
## [11] "LOC121402966" "LOC121398761" "LOC121393708" "LOC121403010" "LOC121399491"
## [16] "LOC121395456" "LOC121393758" "LOC121396893" "LOC121401562" "LOC121401551"
## [21] "LOC108699934" "LOC121400501" "dcc.S"        "LOC121394660" "LOC121396258"
## [26] "LOC108710980" "LOC121400500" "LOC121393376" "LOC121397070" "LOC121402983"
## [31] "c1orf21.S"    "LOC108719816" "LOC108716472" "LOC108698619" "LOC121402182"
## [36] "ada2.L"       "XB22041735.L"
```

```
## [1] "Mapped orthologues"
```

```
## # A tibble: 1 × 3
## # Groups:   cluster [1]
##   cluster       non_Neuronal Neuronal
##   <chr>                <int>    <int>
## 1 Neuroectoderm          187       48
```

### 8 Ascl1 Binding sites (MACS peaks)

```
#Ascl1_peaks_count_xenopus<- load('path/dba_Ascl1_peaks_count_xenopus.RData')

Ascl1_peaks_count_xenopus<- dba(DBAobject, minOverlap = 1) 
Ascl1_GR <- dba.peakset(Ascl1_peaks_count_xenopus, bRetrieve=TRUE)

# Extract the matrix and compute rowMeans
sample_matrix <- as.matrix(mcols(Ascl1_GR)[, c("Ascl1wt_injected_rep1", 
                                               "Ascl1wt_injected_rep2", 
                                               "Ascl1wt_injected_rep3")])
mcols(Ascl1_GR)$Average <- rowMeans(sample_matrix)
print("Ascl1 Chipseq-peaks")
```

```
## [1] "Ascl1 Chipseq-peaks"
```

```
Ascl1_GR
```

```
## GRanges object with 68941 ranges and 4 metadata columns:
##            seqnames              ranges strand | Ascl1wt_injected_rep1
##               <Rle>           <IRanges>  <Rle> |             <numeric>
##       1 NC_054371.1       150224-150824      * |               30.4553
##       2 NC_054371.1       152594-153194      * |              102.1728
##       3 NC_054371.1       189298-189898      * |                0.0000
##       4 NC_054371.1       247379-247979      * |               53.0513
##       5 NC_054371.1       278514-279114      * |                0.0000
##     ...         ...                 ...    ... .                   ...
##   68937 NC_054388.1 116767064-116767664      * |              128.6984
##   68938 NC_054388.1 116907305-116907905      * |              157.1889
##   68939 NC_054388.1 117135750-117136350      * |               59.9283
##   68940 NC_054388.1 117199480-117200080      * |               59.9283
##   68941 NC_054388.1 117217905-117218505      * |               29.4729
##         Ascl1wt_injected_rep2 Ascl1wt_injected_rep3   Average
##                     <numeric>             <numeric> <numeric>
##       1               54.9994               66.9807   50.8118
##       2              126.2950               94.9727  107.8135
##       3               89.6287               49.9856   46.5381
##       4               13.2406               10.9968   25.7629
##       5               32.5923               31.9908   21.5277
##     ...                   ...                   ...       ...
##   68937                0.0000              107.9689   78.8891
##   68938              102.8693              196.9433  152.3338
##   68939              264.8121              162.9531  162.5645
##   68940               96.7583                7.9977   54.8947
##   68941               29.5367               44.9870   34.6656
##   -------
##   seqinfo: 18 sequences from an unspecified genome; no seqlengths
```

```
############################################  filter Ascl1 peaks for peaks with low counts
Ascl1_filtered <- Ascl1_GR[Ascl1_GR$Average >= 100]

# Convert GTF to TxDb
txdb <- makeTxDbFromGFF(gtf_file)

# Annotate peaks
peak_annot <- annotatePeak(
  Ascl1_filtered,
  TxDb = txdb,
  tssRegion = c(-10000, 10000),
  annoDb = NULL  # no orgDB available for Xenopus
)
```

```
## >> preparing features information...      2026-07-07 14:30:33 
## >> identifying nearest features...        2026-07-07 14:30:33 
## >> calculating distance from peak to TSS...   2026-07-07 14:30:34 
## >> assigning genomic annotation...        2026-07-07 14:30:34 
## >> assigning chromosome lengths           2026-07-07 14:30:42 
## >> done...                    2026-07-07 14:30:42
```

```
#..........................................    Figure 1 Study of Ascl1 directed reprograming in Xenopus development. 
plotAnnoBar <- plotAnnoBar(peak_annot)   # this is a ggplot object
#plotAnnoBar

my_cols <- c(
  "Promoter (<=1kb)" = "lightblue",
  "Promoter (1-2kb)" = "#1f78b4",
  "Promoter (2-3kb)" = "#b2df8a",
  "Promoter (3-4kb)" = "#33a02c",
  "Promoter (4-5kb)" = "#fb9a99",
  "Promoter (5-6kb)" = "#e31a1c",
  "Promoter (6-7kb)" = "#fdbf6f",
  "Promoter (7-8kb)" = "#ff7f00",
  "Promoter (8-9kb)" = "#cab2d6",
  "Promoter (9-10kb)"= "#6a3d9a",
  "5' UTR"            = "#ffff99",
  "3' UTR"            = "#b15928",
  "1st Exon"          = "cyan",  # exon colours
  "Other Exon"        = "cyan3",
  "1st Intron"        = "#009966",  # intron colours
  "Other Intron"      = "#006633",
  "Downstream (<=300)"= "grey",
  "Distal Intergenic" = "grey20"
)


plotAnnoBar + scale_fill_manual(values = my_cols)
```

```
txdb<-NULL
peak_annot<-NULL
plotAnnoBar<-NULL
```

#### 8.1 Ascl1 Binding sites regionMatrix to visualise chromatin accessibility: inducible and non-inducible genes scATAC heatmaps

```
## GRanges object with 6 ranges and 6 metadata columns:
##         seqnames            ranges strand | Ascl1wt_injected_rep1
##            <Rle>         <IRanges>  <Rle> |             <numeric>
##   92 NC_054371.1 11076184-11076784      * |               64.8404
##   93 NC_054371.1 11078934-11079534      * |              391.0074
##   94 NC_054371.1 11083162-11083762      * |              366.4466
##   95 NC_054371.1 11106137-11106737      * |              200.4158
##   96 NC_054371.1 11111900-11112500      * |               99.2255
##   97 NC_054371.1 11114863-11115463      * |              111.9971
##      Ascl1wt_injected_rep2 Ascl1wt_injected_rep3   Average   gene_name
##                  <numeric>             <numeric> <numeric> <character>
##   92               93.7027               190.945   116.496    cdc25b.L
##   93              382.9590               611.824   461.930    cdc25b.L
##   94              503.1429               831.760   567.117    cdc25b.L
##   95               81.4806               196.943   159.613    cdc25b.L
##   96              114.0729               227.934   147.078    cdc25b.L
##   97               63.1475               154.955   110.033    cdc25b.L
##      response_class
##         <character>
##   92    InducibleT1
##   93    InducibleT1
##   94    InducibleT1
##   95    InducibleT1
##   96    InducibleT1
##   97    InducibleT1
##   -------
##   seqinfo: 18 sequences from an unspecified genome; no seqlengths
```

#### 8.2 Ascl1 Binding sites boxplots to compare chromatin accessibility

### 9 Motif and chromatin accessibility analysis to identifies potential signatures associated with Ascl1 gene regulation

#### 9.1 HOMER Analysis

```
## [1] "Hommer analysis of NE T1 genes lists"
```

```
## [1] "Top motifs for each class"
```

```
## # A tibble: 47 × 4
##    MotifClass MotifName T1_Neuroectoderm TFs_in_class                           
##    <chr>      <chr>                <dbl> <chr>                                  
##  1 bHLH       Ascl1                  169 Ap4, Ascl1, Ascl2, Atoh1, Atoh7, BHLHA…
##  2 Zf         Slug                    52 Gata2, Gata4, GFY-Staf, GLIS3, HIC1, K…
##  3 Unknown    Tal1                    45 E-box/Drosophila-Promoters/Homer, GAGA…
##  4 bZIP       bZIP52                  33 BATF, bZIP18, bZIP3, bZIP52, bZIP69, C…
##  5 HMG        Sox21                   26 LEF1, SOX1, Sox10, Sox15, Sox17, Sox2,…
##  6 ARF        ARF2                    11 ARF2                                   
##  7 Homeobox   Lhx3                    10 ATHB13, bcd, caudal, Cdx2, CDX4, CRX, …
##  8 MAD        Smad3                   10 Smad2, Smad3, Smad4                    
##  9 Forkhead   Foxo3                    8 FOXA1, Foxa2, Foxa3, Foxf1, FOXK1, Fox…
## 10 T-box      Tbx5                     8 Eomes, Tbet, Tbr1, Tbx21, Tbx5, Tbx6   
## # ℹ 37 more rows
```

#### 9.2 chromVAR

##### 9.2.1 scATAC using Ascl1 Chipseq. peaks of selected list of response genes

```
# keep only larger cell types
cell_types<-levels(multiome_obj@meta.data$cell_type) 
cell_types<-  cell_types[c(-8, -9, -10)]
multiome_obj<- subset(multiome_obj, multiome_obj$cell_type %in% cell_types) 

# get the list of selected response genes from neuroactoderm T1 and T2 and then retrieve chip seq filtered peaks that overlap in 50kb up and intronic regions of permissive genes (sign Up from Neuroactoderm treated vs Neuroactoderm WT)

DefaultAssay(multiome_obj)<-"ATAC"

atac_counts <- FeatureMatrix(fragments = Fragments(multiome_obj), features = selected_genes_peaks, cells = Cells(multiome_obj))

#atac_counts <- counts[, Cells(multiome_obj)]
#print("head(colnames(atac_counts))")
#head(colnames(atac_counts))
#print("head(Cells(multiome_obj))")
#head(Cells(multiome_obj))


multiome_obj@meta.data$ChipATAC_barcodes<- colnames(atac_counts)
multiome_obj[['ChipATAC']] <- CreateChromatinAssay(
  counts = atac_counts,
  sep = c(":", "-"),
  fragments = multiome_obj@assays[["ATAC"]]@fragments[[1]]@path,
  min.cells = 5,
  min.features = 1,
)

# as we are generating CreateChromatinAssay applying filters some values can be NA for cells for that peaks are failing it will not remove cells from RNA assay so generate NA
length(multiome_obj@meta.data$nCount_ChipATAC[!is.na(multiome_obj@meta.data$nCount_ChipATAC)])
```

```
## [1] 22646
```

```
# subset again RNA for CreateChromatinAssay cells that pass  min.cells and min.features filter 
DefaultAssay(multiome_obj)<-"RNA"
multiome_obj <- subset(multiome_obj, cells = rownames(multiome_obj@meta.data)[!is.na(multiome_obj@meta.data$nCount_ChipATAC)])
# again assign the CreateChromatinAssay making sure now cells are common in both RNA and ATAC

DefaultAssay(multiome_obj)<-"ChipATAC"

print("Summary of RNA+ChipATAC object")
```

```
## [1] "Summary of RNA+ChipATAC object"
```

```
multiome_obj
```

```
## An object of class Seurat 
## 521644 features across 22646 samples within 4 assays 
## Active assay: ChipATAC (1322 features, 0 variable features)
##  2 layers present: counts, data
##  3 other assays present: RNA, ATAC, SCT
##  5 dimensional reductions calculated: pca, lsi, wnn.umap, umap.rna, umap.atac
```

```
head(multiome_obj@meta.data)
```

```
##             orig.ident nCount_RNA nFeature_RNA sampleID sample_name
## 01:02:10:04    DNAA001       2799         1598       04    AsclT0r2
## 01:02:45:09    DNAA001        842          676       09   GFPT2_5r1
## 01:02:50:07    DNAA001       1917         1490       07 AsclT1_25r1
## 01:02:73:08    DNAA001        786          570       08 AsclT1_25r2
## 01:02:83:06    DNAA001       1505         1130       06  GFPT1_25r2
## 01:02:90:12    DNAA001        836          640       12  AsclT2_5r2
##             sample_condition     lib Dataset numID percent.mito nCount_ATAC
## 01:02:10:04        +Ascl1_T0 DNAA001  Batch1     2   0.03572705        1104
## 01:02:45:09          WT_T2_5 DNAA001  Batch1     6   0.35629454        1116
## 01:02:50:07     +Ascl1_T1_25 DNAA001  Batch1     9   0.00000000        1920
## 01:02:73:08     +Ascl1_T1_25 DNAA001  Batch1    14   0.00000000         781
## 01:02:83:06         WT_T1_25 DNAA001  Batch1    16   0.13289037         980
## 01:02:90:12      +Ascl1_T2_5 DNAA001  Batch1    18   0.47846890         911
##             nFeature_ATAC frequency_count      FRiP TSS.enrichment
## 01:02:10:04           782            1354 0.8153619       5.494505
## 01:02:45:09           815            1420 0.7859155       6.127206
## 01:02:50:07          1348            2259 0.8499336       4.266004
## 01:02:73:08           574            1051 0.7431018       2.972028
## 01:02:83:06           687            1207 0.8119304       4.368964
## 01:02:90:12           582            1048 0.8692748       3.008103
##             TSS.percentile nCount_SCT nFeature_SCT SCT.weight ATAC.weight
## 01:02:10:04           0.77       2581         1567  0.7688777   0.2311223
## 01:02:45:09           0.84       1454          663  0.7085016   0.2914984
## 01:02:50:07           0.54       1893         1457  0.5133205   0.4866795
## 01:02:73:08           0.23       1409          562  0.7009703   0.2990297
## 01:02:83:06           0.57       1697         1108  0.5584565   0.4415435
## 01:02:90:12           0.24       1476          633  0.7949458   0.2050542
##             wsnn_res.0.1 seurat_clusters cell_type condition  time
## 01:02:10:04            2               2  Mesoderm     ASCL1    T0
## 01:02:45:09            3               3  Endoderm        WT  T2_5
## 01:02:50:07            2               2  Mesoderm     ASCL1 T1_25
## 01:02:73:08            3               3  Endoderm     ASCL1 T1_25
## 01:02:83:06            2               2  Mesoderm        WT T1_25
## 01:02:90:12            2               2  Mesoderm     ASCL1  T2_5
##             ChipATAC_barcodes nCount_ChipATAC nFeature_ChipATAC
## 01:02:10:04       01:02:10:04               2                 2
## 01:02:45:09       01:02:45:09               4                 4
## 01:02:50:07       01:02:50:07              16                13
## 01:02:73:08       01:02:73:08               3                 3
## 01:02:83:06       01:02:83:06               6                 6
## 01:02:90:12       01:02:90:12               1                 1
```

```
########### compute and add ATAC metdadata columns by making sure barcodes consistency
total_counts <- CountFragments(multiome_obj@assays[["ATAC"]]@fragments[[1]]@path)
# Subset total_counts to include only common barcodes
total_counts_subset <- total_counts[total_counts$CB %in% Cells(multiome_obj), ]
#head(total_counts_subset)
#nrow(total_counts_subset)

# Match and add the frequency_count as metadata 
multiome_obj$frequency_count_ChipATAC <- total_counts_subset$frequency_count[match(
  rownames(multiome_obj@meta.data),
  total_counts_subset$CB
)]

Annotation(multiome_obj) <- protein_coding_genes
multiome_obj[['ChipATAC']]
```

```
## ChromatinAssay data with 1322 features for 22646 cells
## Variable features: 0 
## Genome: 
## Annotation present: TRUE 
## Motifs present: FALSE 
## Fragment files: 1
```

```
DefaultAssay(multiome_obj) <- "ChipATAC"

multiome_obj <- RunTFIDF(multiome_obj, assay = "ChipATAC")
multiome_obj <- FindTopFeatures(multiome_obj, min.cutoff = 'q0', assay = "ChipATAC") # 'q0' # Find most frequently observed features
multiome_obj <- RunSVD(multiome_obj,  n = 50, assay = "ChipATAC")
```

##### 9.2.2 Compute motifs deviations scores in ChromVAR assay

```
obj <- multiome_obj
DefaultAssay(obj) <- "ChipATAC"

## 1) Get peak GRanges (already stored correctly in the assay)
gr <- granges(obj[["ChipATAC"]])   # uses proper seqlevels (e.g., NC_054371.1)


## 2) Pull sequences from FASTA file
#fa <- FaFile("........PATH/GCF_017654675.1_Xenopus_laevis_v10.1_genomic.fna")  

 

open(fa)
peak_seqs <- Biostrings::getSeq(fa, gr)
close(fa)

########################################################################### 

## CRUCIAL: force sequence names to EXACTLY match ATAC feature names
names(peak_seqs) <- rownames(obj[["ChipATAC"]])

## 3) Small, focused PFM set (keeps scanning fast)
pfm_all <- getMatrixSet(JASPAR2022, list(collection="CORE", tax_group="vertebrates"))
# Make motif names unique (avoid the warning and ensure rownames match)
nm <- sapply(pfm_all, name); names(pfm_all) <- make.unique(nm)

############################################# 

## 4) Compute BOTH matches (binary matrix) and positions
#mm_matches <- motifmatchr::matchMotifs(
#  pwms   = pfm_all,
 # subject  = peak_seqs,
#  out      = "matches",
#  p.cutoff = 5e-4
#)
#motif_ix <- motifmatchr::motifMatches(mm_matches)     # sparse matrix (motifs x peaks)

 #motif_pos <- motifmatchr::matchMotifs( 
 # pwms   = pfm_all,
 # subject  = peak_seqs,
 # out      = "positions",
 # p.cutoff = 5e-4
 #)
 
# 5) Add row/col names to the matches matrix so Signac can align to features
#motif_ix2 <- as(motif_ix, "dMatrix")    # coerce to numeric sparse
#rownames(motif_ix2) <- rownames(obj[["ChipATAC"]])  # peaks (features)
# Reorder columns to match pfm names exactly
#motif_ix2 <- motif_ix2[, names(pfm_all), drop = FALSE]

# 6) Sanity checks (important)
#stopifnot(names(peak_seqs) == rownames(obj[["ChipATAC"]]) )
#stopifnot(nrow(motif_ix2) == nrow(obj[["ChipATAC"]]))
#stopifnot(identical(colnames(motif_ix2), names(pfm_all)))
#stopifnot(length(motif_pos) == length(pfm_all))
#stopifnot(all(names(motif_pos) %in% names(pfm_all)))

# (optional) reorder positions list to pfm order
#motif_pos <- motif_pos[names(pfm_all)]

# 7) Build and attach the Motif object
#motobj <- CreateMotifObject(
#  data      = motif_ix2,   # peaks x motifs (rownames = ATAC features)
#  pwm       = pfm_all,
#  positions = motif_pos
#)

#obj <- SetAssayData(obj, assay = "ChipATAC", slot = "motifs", new.data = motobj)

# 9) Verify it worked
#Motifs(obj)            # should list 3 motifs, not 0


# 10) chromVAR on cells
#obj <- RunTFIDF(obj)
#obj <- RunChromVAR(obj, genome = fa) # it takes about 200 GB RAM 


obj <-readRDS(paste0(obj_path, "chromvar_mesodermT12.rds"))  # to save computing time run once and save object to read quickly here

print("chromVAR assay now exists")
```

```
## [1] "chromVAR assay now exists"
```

```
# sanity check: chromVAR assay now exists
obj                      # should include "chromvar"
```

```
## An object of class Seurat 
## 522485 features across 22647 samples within 5 assays 
## Active assay: chromvar (841 features, 0 variable features)
##  1 layer present: data
##  4 other assays present: RNA, ATAC, SCT, ChipATAC
##  5 dimensional reductions calculated: pca, lsi, wnn.umap, umap.rna, umap.atac
```

```
# Visualize
DefaultAssay(obj) <- "chromvar"
# Check available motif names:
#rownames(GetAssayData(obj, "chromvar"))  # e.g., "ASCL1","ASCL1.1","ZEB1"

# Parse condition/time once
obj$condition <- ifelse(grepl("^\\+?Ascl1", obj$sample_condition), "ASCL1", "WT")
obj$time      <- sub("^(\\+?Ascl1|WT)_", "", obj$sample_condition)
```

##### 9.2.3 Find differentially accessible motifs between cell types (+Ascl1\_T1\_25 and +Ascl1\_T2\_5)

List of comparisons (Neuroectoderm vs other cell types one by
one)

- Neuroectoderm vs Endoderm
- Neuroectoderm vs Mesoderm
- Neuroectoderm vs Non neuroectoderm
- Neuroectoderm vs Fibroblast
- Neuroectoderm vs Lateral-plate-mesoderm
- Neuroectoderm vs Epithelial-skin

```
#####################################################
#####################################################
#####################################################


obj <- subset(obj, sample_condition %in% c("+Ascl1_T1_25", "+Ascl1_T2_5"))


DefaultAssay(obj) <- "chromvar"
obj
```

```
## An object of class Seurat 
## 522485 features across 8474 samples within 5 assays 
## Active assay: chromvar (841 features, 0 variable features)
##  1 layer present: data
##  4 other assays present: RNA, ATAC, SCT, ChipATAC
##  5 dimensional reductions calculated: pca, lsi, wnn.umap, umap.rna, umap.atac
```

```
print("+Ascl1 T1 and T2: Cell count summary for DA analysis")
```

```
## [1] "+Ascl1 T1 and T2: Cell count summary for DA analysis"
```

```
table(obj$cell_type)
```

```
## 
##               Mesoderm          Neuroectoderm               Endoderm 
##                   1919                   2258                   1743 
## Lateral-plate-mesoderm        Epithelial-skin      Non-neuroectoderm 
##                    828                    734                    746 
##             Fibroblast 
##                    246
```

```
table(obj@meta.data$time[obj@meta.data$condition == "ASCL1"])
```

```
## 
## T1_25  T2_5 
##  4953  3521
```

```
mat <- GetAssayData(obj, slot = "data")  # motifs x cells
#head(t(mat))
# Option A: keep only motifs with ZERO NAs
#keep_motifs <- rowSums(is.na(mat)) == 0

# Option B (softer): keep motifs with <=10% NAs
keep_motifs <- rowMeans(is.na(mat)) <= 0.20

#sum(keep_motifs)  # how many motifs survive?

# Subset the chromVAR assay to those motifs
obj[["chromvar"]] <- subset(obj[["chromvar"]],
                                    features = rownames(mat)[keep_motifs])

#####################
# 1) Pull the chromVAR deviation matrix (motifs x cells)
mat <- GetAssayData(obj[["chromvar"]], slot = "data")

# 2) Identify cells with ZERO NAs
keep_cells_lgl <- Matrix::colSums(is.na(mat)) == 0
keep_cells <- colnames(mat)[keep_cells_lgl]

cat("Cells kept after removing cells with NAs:", length(keep_cells), "of", ncol(mat), "\n")
```

```
## Cells kept after removing cells with NAs: 7095 of 8474
```

```
# 3) Subset the whole Seurat object to those cells
# (this will subset all assays/metadata consistently)
obj <- subset(obj, cells = keep_cells)

# 4) (Optional) sanity check: chromVAR matrix now has no NAs
mat2 <- GetAssayData(obj[["chromvar"]], slot = "data")
stopifnot(all(Matrix::colSums(is.na(mat2)) == 0))

#print("subset for only +Ascl1_T1_25")
#head(obj@meta.data)
cell_types <- unique(obj$cell_type)  # Get unique cell types

##############################################################################
################################################### 
##############..................

Idents(obj)<-obj$cell_type


# List of comparisons (Neuroectoderm vs other cell types)
comparisons <- list(
  c("Neuroectoderm","Endoderm" ) ,
  c("Neuroectoderm","Mesoderm" ),
  c("Neuroectoderm", "Non-neuroectoderm"),
  c("Neuroectoderm", "Lateral-plate-mesoderm"),
  c("Neuroectoderm", "Epithelial-skin") )

#########################################################
######## DA Motifs analysis between Neuroectoderm vs all celltypes one by one for all Motifs (ChromVar assay)
#######################################################
print("DA Motifs analysis between Neuroectoderm vs all celltypes one by one for all Motifs")
```

```
## [1] "DA Motifs analysis between Neuroectoderm vs all celltypes one by one for all Motifs"
```

```
# Initialize an empty list to store results
all_da_motifs <- list()

# Loop through each comparison pair
for (comp in comparisons) {
  ident1 <- comp[1]
  ident2 <- comp[2]
  
  # Run differential accessibility (DA) analysis
  da_motifs <- FindMarkers(
    object = obj,
    assay = "chromvar",
    ident.1 = ident1,
    ident.2 = ident2,
    test.use = 'LR',  #  "LR"
    min.pct = 0.1,
    group.by = 'cell_type',
    #latent.vars = "nCount_ATAC"
  )

  
  # Add peak names
  da_motifs$motifs <- rownames(da_motifs)
  rownames(da_motifs) <- NULL  # Reset row names

  # Add a column for the comparison name
  da_motifs$Comparison <- paste(ident1, "vs", ident2)

  # Reorder columns
  da_motifs <- da_motifs[, c("motifs", "Comparison", "p_val_adj", "p_val", "avg_log2FC", "pct.1", "pct.2")]

  # Store in list
  all_da_motifs[[paste(ident1, ident2, sep = "_vs_")]] <- da_motifs
}

# Combine all results into a single data frame
final_da_table <- do.call(rbind, all_da_motifs)

# View the final combined table
final_da_table <- final_da_table %>%
  filter(p_val < 0.05)
print("all results after filtering p_val < 0.05 ")
```

```
## [1] "all results after filtering p_val < 0.05 "
```

```
datatable(final_da_table, 
          class = "compact",
          filter = "top",
          rownames = FALSE,
          colnames = c("Motif", "Comparison", "Adj. P-value", "P-value", "Avg. Log2FC", 
                       "% Cells Accessible in Group 1", "% Cells Accessible in Group 2"),
          extensions = c('Buttons'),
          options = list(
            pageLength = 15,
            dom = 'Bfrtip',
            buttons = c('excel', 'csv', 'pdf', 'copy')
          ))
```

##### 9.2.4 Top Motifs from HOMER and chromVAR

```
##  [1] "ASCL1"  "Ascl2"  "MYOG"   "ZBTB18" "ZEB1"   "TGIF2"  "TBX5"   "Foxo3" 
##  [9] "Foxf1"  "FOXP1"  "FOXK1"  "EOMES"  "TBR1"   "TBX21"
```

### 10 Sox motif family expression data visualization

### 11 Session information

```
sessionInfo()
```

```
## R version 4.4.1 (2024-06-14)
## Platform: x86_64-pc-linux-gnu
## Running under: Ubuntu 22.04.4 LTS
## 
## Matrix products: default
## BLAS:   /usr/lib/x86_64-linux-gnu/openblas-pthread/libblas.so.3 
## LAPACK: /usr/lib/x86_64-linux-gnu/openblas-pthread/libopenblasp-r0.3.20.so;  LAPACK version 3.10.0
## 
## locale:
##  [1] LC_CTYPE=en_GB.UTF-8       LC_NUMERIC=C              
##  [3] LC_TIME=en_GB.UTF-8        LC_COLLATE=en_GB.UTF-8    
##  [5] LC_MONETARY=en_GB.UTF-8    LC_MESSAGES=en_GB.UTF-8   
##  [7] LC_PAPER=en_GB.UTF-8       LC_NAME=C                 
##  [9] LC_ADDRESS=C               LC_TELEPHONE=C            
## [11] LC_MEASUREMENT=en_GB.UTF-8 LC_IDENTIFICATION=C       
## 
## time zone: Europe/London
## tzcode source: system (glibc)
## 
## attached base packages:
## [1] stats4    stats     graphics  grDevices utils     datasets  methods  
## [8] base     
## 
## other attached packages:
##  [1] tidytext_0.4.3              enrichplot_1.26.6          
##  [3] org.Mm.eg.db_3.20.0         org.Hs.eg.db_3.20.0        
##  [5] clusterProfiler_4.14.6      tibble_3.3.0               
##  [7] ggtext_0.1.2                JASPAR2022_0.99.8          
##  [9] BiocFileCache_2.14.0        dbplyr_2.5.0               
## [11] TFBSTools_1.44.0            Rsamtools_2.22.0           
## [13] Biostrings_2.74.1           XVector_0.46.0             
## [15] cowplot_1.2.0               ggseqlogo_0.2              
## [17] universalmotif_1.24.2       pheatmap_1.0.13            
## [19] purrr_1.1.0                 tidyr_1.3.1                
## [21] rvest_1.0.4                 GenomicFeatures_1.58.0     
## [23] AnnotationDbi_1.68.0        ChIPseeker_1.42.1          
## [25] DiffBind_3.16.0             profileplyr_1.22.0         
## [27] SummarizedExperiment_1.36.0 Biobase_2.66.0             
## [29] MatrixGenerics_1.18.1       matrixStats_1.5.0          
## [31] DT_0.33                     future_1.67.0              
## [33] Matrix_1.7-3                rtracklayer_1.66.0         
## [35] GenomicRanges_1.58.0        GenomeInfoDb_1.42.3        
## [37] IRanges_2.40.1              S4Vectors_0.44.0           
## [39] BiocGenerics_0.52.0         patchwork_1.3.1            
## [41] stringr_1.5.1               dplyr_1.1.4                
## [43] ggplot2_3.5.2               Signac_1.14.0              
## [45] Seurat_5.3.0                SeuratObject_5.1.0         
## [47] sp_2.2-0                   
## 
## loaded via a namespace (and not attached):
##   [1] R.methodsS3_1.8.2                        
##   [2] dichromat_2.0-0.1                        
##   [3] progress_1.2.3                           
##   [4] tiff_0.1-12                              
##   [5] poweRlaw_1.0.0                           
##   [6] goftest_1.2-3                            
##   [7] TxDb.Hsapiens.UCSC.hg38.knownGene_3.20.0 
##   [8] vctrs_0.6.5                              
##   [9] ggtangle_0.0.7                           
##  [10] spatstat.random_3.4-1                    
##  [11] digest_0.6.37                            
##  [12] png_0.1-8                                
##  [13] shape_1.4.6.1                            
##  [14] ggrepel_0.9.6                            
##  [15] mixsqp_0.3-54                            
##  [16] deldir_2.0-4                             
##  [17] parallelly_1.45.1                        
##  [18] MASS_7.3-61                              
##  [19] reshape2_1.4.4                           
##  [20] SQUAREM_2021.1                           
##  [21] httpuv_1.6.16                            
##  [22] foreach_1.5.2                            
##  [23] qvalue_2.38.0                            
##  [24] withr_3.0.2                              
##  [25] xfun_0.52                                
##  [26] amap_0.8-20                              
##  [27] ggfun_0.2.0                              
##  [28] survival_3.7-0                           
##  [29] commonmark_2.0.0                         
##  [30] memoise_2.0.1                            
##  [31] gson_0.1.0                               
##  [32] tidytree_0.4.6                           
##  [33] zoo_1.8-14                               
##  [34] GlobalOptions_0.1.2                      
##  [35] gtools_3.9.5                             
##  [36] pbapply_1.7-4                            
##  [37] R.oo_1.27.1                              
##  [38] prettyunits_1.2.0                        
##  [39] KEGGREST_1.46.0                          
##  [40] promises_1.3.3                           
##  [41] httr_1.4.7                               
##  [42] GreyListChIP_1.38.0                      
##  [43] restfulr_0.0.16                          
##  [44] globals_0.18.0                           
##  [45] fitdistrplus_1.2-4                       
##  [46] ashr_2.2-63                              
##  [47] rstudioapi_0.17.1                        
##  [48] UCSC.utils_1.2.0                         
##  [49] miniUI_0.1.2                             
##  [50] generics_0.1.4                           
##  [51] DOSE_4.0.1                               
##  [52] curl_6.4.0                               
##  [53] zlibbioc_1.52.0                          
##  [54] EnrichedHeatmap_1.36.0                   
##  [55] polyclip_1.10-7                          
##  [56] GenomeInfoDbData_1.2.13                  
##  [57] SparseArray_1.6.2                        
##  [58] xtable_1.8-4                             
##  [59] doParallel_1.0.17                        
##  [60] evaluate_1.0.4                           
##  [61] S4Arrays_1.6.0                           
##  [62] systemPipeR_2.12.0                       
##  [63] preprocessCore_1.68.0                    
##  [64] hms_1.1.3                                
##  [65] irlba_2.3.5.1                            
##  [66] filelock_1.0.3                           
##  [67] colorspace_2.1-1                         
##  [68] ROCR_1.0-11                              
##  [69] reticulate_1.43.0                        
##  [70] spatstat.data_3.1-6                      
##  [71] readr_2.1.5                              
##  [72] magrittr_2.0.3                           
##  [73] lmtest_0.9-40                            
##  [74] later_1.4.2                              
##  [75] ggtree_3.14.0                            
##  [76] lattice_0.22-7                           
##  [77] spatstat.geom_3.5-0                      
##  [78] future.apply_1.20.0                      
##  [79] scattermore_1.2                          
##  [80] XML_3.99-0.18                            
##  [81] RcppAnnoy_0.0.22                         
##  [82] pillar_1.11.0                            
##  [83] nlme_3.1-168                             
##  [84] iterators_1.0.14                         
##  [85] pwalign_1.2.0                            
##  [86] caTools_1.18.3                           
##  [87] compiler_4.4.1                           
##  [88] RSpectra_0.16-2                          
##  [89] stringi_1.8.7                            
##  [90] tokenizers_0.3.0                         
##  [91] tensor_1.5                               
##  [92] GenomicAlignments_1.42.0                 
##  [93] plyr_1.8.9                               
##  [94] crayon_1.5.3                             
##  [95] abind_1.4-8                              
##  [96] BiocIO_1.16.0                            
##  [97] truncnorm_1.0-9                          
##  [98] gridGraphics_0.5-1                       
##  [99] emdbook_1.3.14                           
## [100] locfit_1.5-9.12                          
## [101] bit_4.6.0                                
## [102] fastmatch_1.1-6                          
## [103] codetools_0.2-20                         
## [104] TxDb.Mmusculus.UCSC.mm10.knownGene_3.10.0
## [105] crosstalk_1.2.1                          
## [106] bslib_0.9.0                              
## [107] TxDb.Hsapiens.UCSC.hg19.knownGene_3.2.2  
## [108] GetoptLong_1.0.5                         
## [109] plotly_4.11.0                            
## [110] leidenbase_0.1.35                        
## [111] mime_0.12                                
## [112] splines_4.4.1                            
## [113] markdown_2.0                             
## [114] circlize_0.4.16                          
## [115] Rcpp_1.1.0                               
## [116] fastDummies_1.7.5                        
## [117] interp_1.1-6                             
## [118] utf8_1.2.6                               
## [119] gridtext_0.1.6                           
## [120] knitr_1.50                               
## [121] blob_1.2.4                               
## [122] seqLogo_1.72.0                           
## [123] clue_0.3-66                              
## [124] apeglm_1.28.0                            
## [125] chipseq_1.56.0                           
## [126] fs_1.6.6                                 
## [127] listenv_0.9.1                            
## [128] ggplotify_0.1.2                          
## [129] rGREAT_2.8.0                             
## [130] statmod_1.5.0                            
## [131] tzdb_0.5.0                               
## [132] pkgconfig_2.0.3                          
## [133] tools_4.4.1                              
## [134] cachem_1.1.0                             
## [135] RSQLite_2.4.2                            
## [136] viridisLite_0.4.2                        
## [137] DBI_1.2.3                                
## [138] numDeriv_2016.8-1.1                      
## [139] fastmap_1.2.0                            
## [140] rmarkdown_2.29                           
## [141] scales_1.4.0                             
## [142] grid_4.4.1                               
## [143] ica_1.0-3                                
## [144] sass_0.4.10                              
## [145] coda_0.19-4.1                            
## [146] dotCall64_1.2                            
## [147] selectr_0.4-2                            
## [148] RANN_2.6.2                               
## [149] farver_2.1.2                             
## [150] yaml_2.3.10                              
## [151] latticeExtra_0.6-30                      
## [152] cli_3.6.5                                
## [153] txdbmaker_1.2.1                          
## [154] lifecycle_1.0.4                          
## [155] uwot_0.2.3                               
## [156] mvtnorm_1.3-3                            
## [157] annotate_1.84.0                          
## [158] BiocParallel_1.40.2                      
## [159] gtable_0.3.6                             
## [160] rjson_0.2.23                             
## [161] ggridges_0.5.6                           
## [162] progressr_0.15.1                         
## [163] SnowballC_0.7.1                          
## [164] parallel_4.4.1                           
## [165] ape_5.8-1                                
## [166] limma_3.62.2                             
## [167] jsonlite_2.0.0                           
## [168] RcppHNSW_0.6.0                           
## [169] bitops_1.0-9                             
## [170] bit64_4.6.0-1                            
## [171] Rtsne_0.17                               
## [172] yulab.utils_0.2.0                        
## [173] litedown_0.7                             
## [174] spatstat.utils_3.1-5                     
## [175] soGGi_1.38.0                             
## [176] CNEr_1.42.0                              
## [177] janeaustenr_1.0.0                        
## [178] bdsmatrix_1.3-7                          
## [179] jquerylib_0.1.4                          
## [180] GOSemSim_2.32.0                          
## [181] spatstat.univar_3.1-4                    
## [182] R.utils_2.13.0                           
## [183] lazyeval_0.2.2                           
## [184] shiny_1.11.1                             
## [185] htmltools_0.5.8.1                        
## [186] GO.db_3.20.0                             
## [187] sctransform_0.4.2                        
## [188] rappdirs_0.3.3                           
## [189] glue_1.8.0                               
## [190] TFMPvalue_0.0.9                          
## [191] TxDb.Mmusculus.UCSC.mm9.knownGene_3.2.2  
## [192] spam_2.11-1                              
## [193] httr2_1.2.1                              
## [194] RCurl_1.98-1.17                          
## [195] treeio_1.30.0                            
## [196] BSgenome_1.74.0                          
## [197] jpeg_0.1-11                              
## [198] gridExtra_2.3                            
## [199] boot_1.3-31                              
## [200] igraph_2.1.4                             
## [201] invgamma_1.2                             
## [202] R6_2.5.1                                 
## [203] gplots_3.2.0                             
## [204] labeling_0.4.3                           
## [205] RcppRoll_0.3.1                           
## [206] cluster_2.1.6                            
## [207] bbmle_1.0.25.1                           
## [208] aplot_0.2.8                              
## [209] DirichletMultinomial_1.48.0              
## [210] DelayedArray_0.32.0                      
## [211] tidyselect_1.2.1                         
## [212] plotrix_3.8-4                            
## [213] xml2_1.3.8                               
## [214] KernSmooth_2.23-24                       
## [215] data.table_1.17.8                        
## [216] htmlwidgets_1.6.4                        
## [217] fgsea_1.32.4                             
## [218] ComplexHeatmap_2.22.0                    
## [219] RColorBrewer_1.1-3                       
## [220] hwriter_1.3.2.1                          
## [221] biomaRt_2.62.1                           
## [222] rlang_1.1.6                              
## [223] spatstat.sparse_3.1-0                    
## [224] spatstat.explore_3.5-2                   
## [225] ShortRead_1.64.0
```
